# JointMap identified SERPINA3^+^ chondrocytes as a therapeutic target for osteoarthritis

**DOI:** 10.1101/2025.09.09.675251

**Authors:** Haoda Wu, Wenqiang Yan, Mengze Sun, Yifei Fan, Shaowei Jia, Junyan Wang, Boyang Xu, Boyun Lu, Yuqing Du, Long Chen, Jian Lin, Jin Cheng, Yingfang Ao, Xiaoqing Hu

## Abstract

**Background:** Osteoarthritis (OA) is a prevalent degenerative joint disease that affects nearly half of the population aged over 65. Recent single-cell transcriptional studies have identified heterogeneous OA-specific chondrocyte subtypes, yet a unified subtype-classification remains incomplete.

**Methods:** In this study, we integrated a total of 545,946 cells to develop ‘JointMap’, an interspecies atlas. The chondrocyte subtypes were analyzed using machine learning algorithms were validated via human samples, animal models, and in vitro cell culture. We performed KAS-seq and analyzed the spatial characters for JointMap.

**Results:** Based on clinical diagnosis and cross-omic indexes, we identified a novel OA-elevated chondrocyte subtype, SERPINA3^+^ chondrocytes. Notably, we uncovered their upregulated chondrogenesis function and intercellular communication patterns as potential therapeutic targets.

**Conclusions:** In summary, JointMap provides a unified framework for the systematic annotation of chondral tissues across species and helps in the identification of druggable networks for articular diseases.

## INTRODUCTION

Articular cartilage plays a lifelong role in joint lubrication and load transmission [1–4]. However, its regenerative capacity is limited due to the lack of vasculature, resulting in substantial clinical and economic burdens [1]. Osteoarthritis (OA), the most prevalent form of cartilage degeneration, is predicted to affect 78.4 million individuals in the United States by 2040 [5, 6]. This chronic disabling condition currently impacts approximately 300 million people worldwide, with more than half of adults aged 65 or older exhibiting clinical manifestations [7].

Recent single-cell transcriptional studies have identified heterogeneous chondrocyte subtypes within articular cartilage and associated tissues [2–4, 8–21]. These subtypes exhibit distinct molecular characteristics and play corresponding pathological roles in degenerated cartilage, OA, rheumatoid arthritis (RA), and other conditions [2–4, 8–21]. Therefore, unifying chondrocyte classification is of great significance. Nevertheless, due to the limited amount of data, previously proposed classification standards remain inconsistent across studies. This variation hinders communication among researchers and reduces the universality of the conclusions. However, for standardizing the classification of chondrocytes, a large unified chondral atlas across species remains incompletely constructed [2, 3, 22].

To bridge this gap, we developed ‘JointMap’, an interspecies single-cell atlas incorporating clinical, spatial, and multi-omic annotations: (1) JointMap incorporates 14 chondral datasets encompassing 545,946 cells and 22,161 genes; (2) the 37 chondrocyte subtypes were classified using machine learning algorithms and annotated by multi-omic datasets; (3) the OA-elevated SERPINA3^+^ chondrocytes altered intercellular communication and promoted chondrogenesis. Collectively, as a publicly available platform, JointMap provides researchers with annotated reference datasets that enable exploration in cartilage pathology.

## RESULTS

### The generation of JointMap

To construct a unified chondral atlas encompassing cells from major joint tissues, we integrated 14 publicly available single-cell datasets, focusing on articular cartilage, meniscus, and associated joint tissues (Fig. 1A; Table S1) [2, 4, 8–17]. The datasets were systematically analyzed using the Seurat and Harmony algorithms, with count matrix under normalized rank process to minimize batch effect [20, 23, 24]. After stringent quality control, JointMap comprised 545,946 cells and 22,161 genes (Fig. S1A).

**Figure 1.**
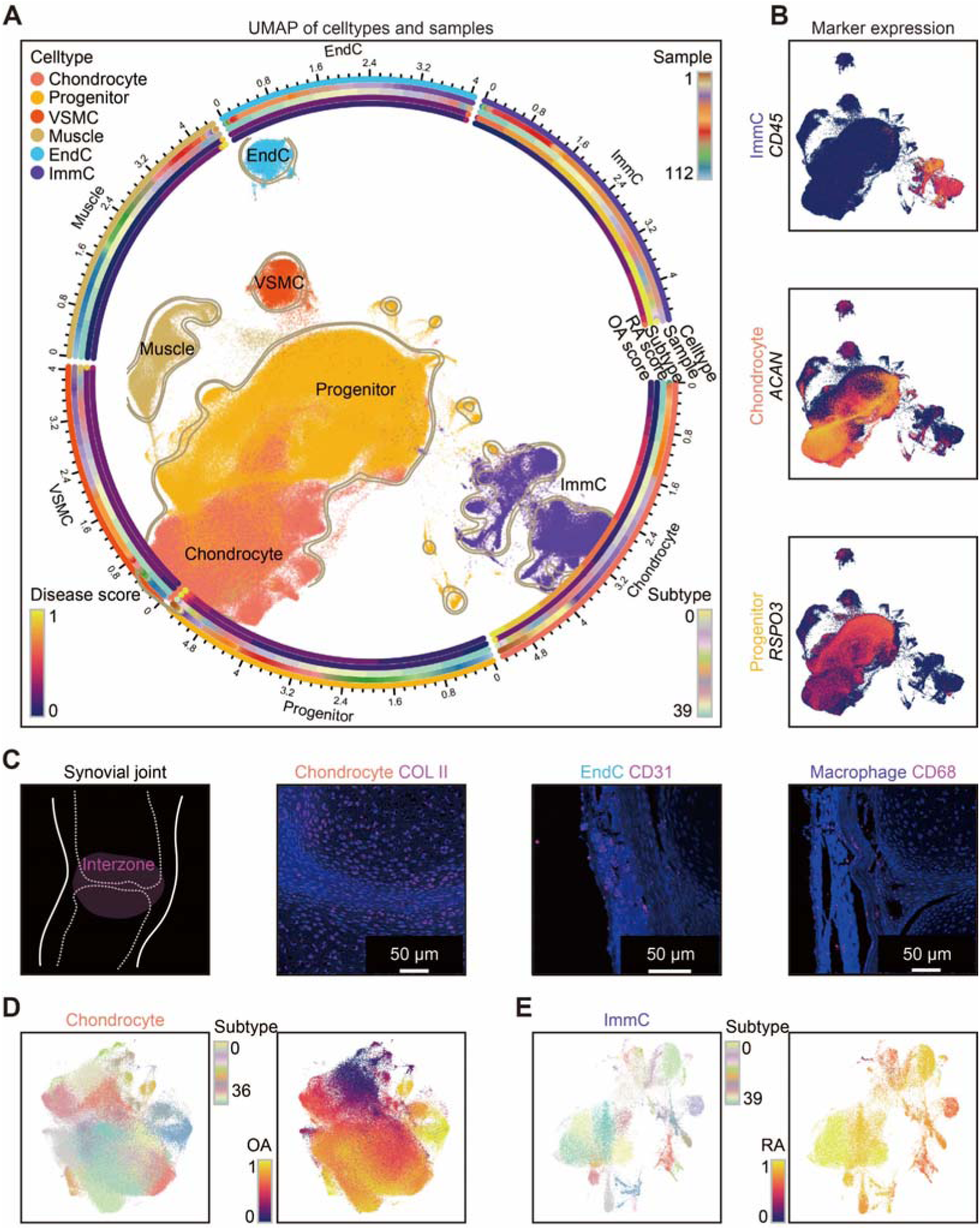
Unified chondral atlas JointMap. (A) Embedding of JointMap showing major cell populations. The circles from the outside to the inside presenting the major cell populations, samples, subtypes, RA scores, and OA scores. (B) Embedding showing the expression of key markers. (C) Anatomical sampling and immunostaining of DAPI (blue) and markers (COL II for chondrocytes, CD31 for EndCs, and CD68 for macrophages; red) in nascent synovial joint of one-week-old mice. (D) Re-embedding of chondrocyte subtypes and their OA scores. (E) Re-embedding of ImmC subtypes and their RA scores.

JointMap contained six major cell populations: chondrocytes (*ACAN*, n = 204,649), progenitors (*RSPO3*, n = 250,150), muscle cells (*MYOG*, n = 25,440), vascular smooth muscle cells (VSMCs, *MYH11*, n = 11,091), endothelial cells (EndCs, *CD31*, n = 11,535), and immune cells (ImmCs, *CD45*, n = 43,081) (Fig. 1B and Fig. S1B). Notably, the continuity observed in the uniform manifold approximation and projection (UMAP) between chondrocytes and progenitors suggested a close lineage [25]. Further stratification using cell-cycle analysis and stemness analysis refined these populations (Fig. S1C) [21, 26]. Collectively, JointMap provided a visualization of the synovial joint cellular composition, offering insights into potential cell-cell interactions (Fig. 1C).

JointMap encompasses diverse disease conditions, including degeneration, regeneration, OA, RA, and *Sox9* deficient, alongside healthy and wildtype controls. This enabled us to calculate disease-specific scores in single-cell level (Fig. 1A and Fig. S1D) [2, 4, 8–17]. Articular cartilage is essential for cushioning and absorbing mechanical forces [1]. However, due to the lack of blood vessels and nerves, its ability for self-repair and regeneration is severely limited [22]. Cartilage degeneration or injuries exceeding 2 cm² often lead to OA [1]. Similarly, RA is a chronic inflammatory autoimmune disorder primarily affecting synovial joints [27]. Genetic variations in human leukocyte antigen genes are strongly associated with RA development [27]. Reflecting these clinical associations, OA scores were markedly elevated in chondrocytes, whereas RA scores were highest in ImmCs, validating the disease-specific annotations in JointMap (Fig. S1D).

To bridge the gap between clinical diagnosis and cell subtypes, the six major cell populations were further visualized and clustered into finer subtypes (Fig. 1D, E and Fig. S1E). Notably, specific chondrocyte subtypes exhibited pronounced OA scores, implicating their potential role in OA progression (Fig. 1D). Similarly, RA scores were significantly elevated in specific ImmC subtypes (Fig. 1E). These associations will be further explored in subsequent research.

### Chondrocyte subtypes and arthritis

Previous studies have identified numerous chondrocyte subtypes, including hyaline cartilage chondrocytes (localized in articular cartilage), fibrocartilage chondrocytes (localized in the menisci), homeostatic chondrocytes, hypertrophic chondrocytes, proliferative chondrocytes, regulatory chondrocytes, effector chondrocytes, resting chondrocytes, aging chondrocytes, growth plate chondrocytes, osteochondrocytes, superficial chondrocytes, deep chondrocytes, lipochondrocyte, and numerically or genetically designated subtypes [2–4, 8–21]. These findings underscore the remarkable heterogeneity of chondrocyte populations. To illustrate this diversity, we compiled a subset of chondrocyte subtypes along with their reported marker genes (Table S2). However, existing classification standards exhibit partial overlap in marker genes, highlighting the need for a more refined approach. To elucidate the molecular profiles of 204,649 chondrocytes within JointMap, we implemented high-resolution gradient clustering for subtype classification (Fig. S2A). These results showed both similarity and novelty to the cell identities annotated in original studies (Fig. S2B-D; Table S1).

We employed the l_2_-norm regularized logistic regression-based machine learning algorithm to infer molecular characteristics of chondrocyte subtypes (Fig. 2A; Table S2) [31, 32]. To further elucidate transcriptional regulatory network, we assessed the regulon activity of transcription factors predicted by SCENIC (Fig. 2A) [33]. Together, these analyses established an objective classification standard for chondrocyte subtypes based on single-cell big data with machine learning methodologies.

**Figure 2.**
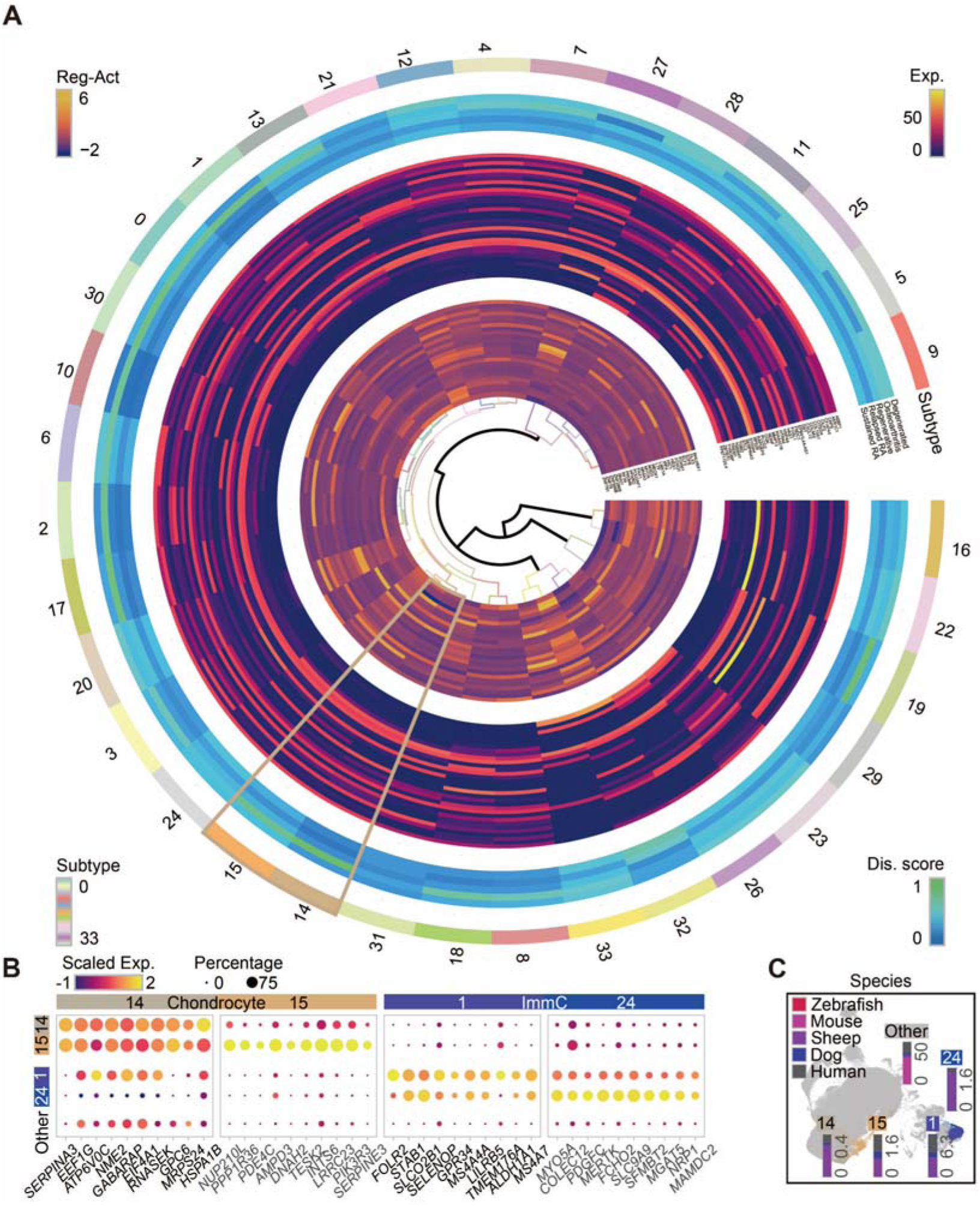
Harmonized annotation of chondrocyte subtypes. (A) Circular heatmaps from the outside to the inside presenting the chondrocyte subtypes, disease (Dis.) scores (including degenerated score, OA score, regenerative score, relapsed RA score, and sustained RA score), gene expression (Exp.) of markers inferred by machine learning (Table S2) [31, 32], and regulons activity (Reg-Act) speculated by SCENIC [33]. (B) Dot plots displaying marker genes of chondrocyte subtype 14/15 and ImmC subtype 1/24. Dot color corresponding to the scaled expression level, and dot size to the percentage of cells expressing a marker. (C) Embedding showing chondrocyte subtype 14/15 and ImmC subtype 1/24. Bars displaying the cell percentage of different species for corresponding subtypes.

Notably, chondrocyte subtype 14/15 exhibited elevated OA score (Fig. 2A). Given that over 50% of individuals aged 65 or older are diagnosed with OA [7], these subtypes displayed a higher composition of elderly samples (*p* = 0.045; Fig. S2E). Machine learning-derived molecular traits revealed distinct expression patterns for chondrocyte subtype 14/15 (Fig. S2F) identifying SERPINA3 as a key marker gene (Fig. 2B). SERPINA3^+^ chondrocytes constituted approximately 6% of the total chondrocyte population (Fig. S2A). Recent genome-wide association study has identified SERPINA3 as one of effector genes involved in OA [34]. Combining together, the elevation of OA scores in SERPINA3^+^ chondrocytes suggest functional significance in OA pathogenesis.

Conservation across species is critical for developing animal models in therapeutic research. Recent studies have implicated SERPINA3 in neocortical folding and cognitive enhancement during brain evolution [35, 36]. Here, we demonstrated that SERPINA3^+^ chondrocytes were not exclusive to humans (Fig. 2C and Fig. S2G), supporting the feasibility for preclinical investigations using animal disease models. Regarding interspecies differences, we analyzed regulon activities across human, dog, sheep, and mouse datasets (Fig. S2H; Table S2). Approximately 200 regulons were exhibited significant interspecies variation within each subtype. Notably, murine chondrocytes exhibited more species-specific regulons. On the contrast, the progenitors and ImmCs exhibited more human-specific regulons. At least 35% of these subtype-specific interspecies regulons were detected in major cell population level. For instance, about 50% of human-specific regulons in chondrocyte subtype 0 were also variated in human chondrocytes compared to chondrocytes from other species. Overall, while cell types and their subtypes exhibited unique interspecies variation patterns, the core regulatory networks remained highly conserved (76%; 700/925).

### Arthritic transcriptional change in chondrocyte subtypes

OA is a progressive and disabling disease, two traits of which are inflammation and the loss of articular cartilage (Fig. 3A) [4, 39, 40]. SERPINA3 was identified as the effector gene of OA based on 1,962,069 individuals [34]. Among the various molecules involved in OA pathogenesis, SERPINA3 has emerged as a significant factor [22, 41, 42]. As an acute-phase response protein, SERPINA3 exhibits anti-inflammatory functions. Specifically, silencing SERPINA3 in human OA primary chondrocytes results in a 22% increase in the expression of the NF-κB inflammatory pathway [22]. Furthermore, elevated serum SERPINA3 level are positively correlated with the severity and duration of inflammation [41, 42]. Notably, upregulation of SERPINA3 using Sinensetin reduces OA-related inflammation and alleviates articular cartilage damage [22]. These findings collectively suggest that SERPINA3 functions as an inhibitor of inflammatory factors and a chondroprotective agent.

**Figure 3.**
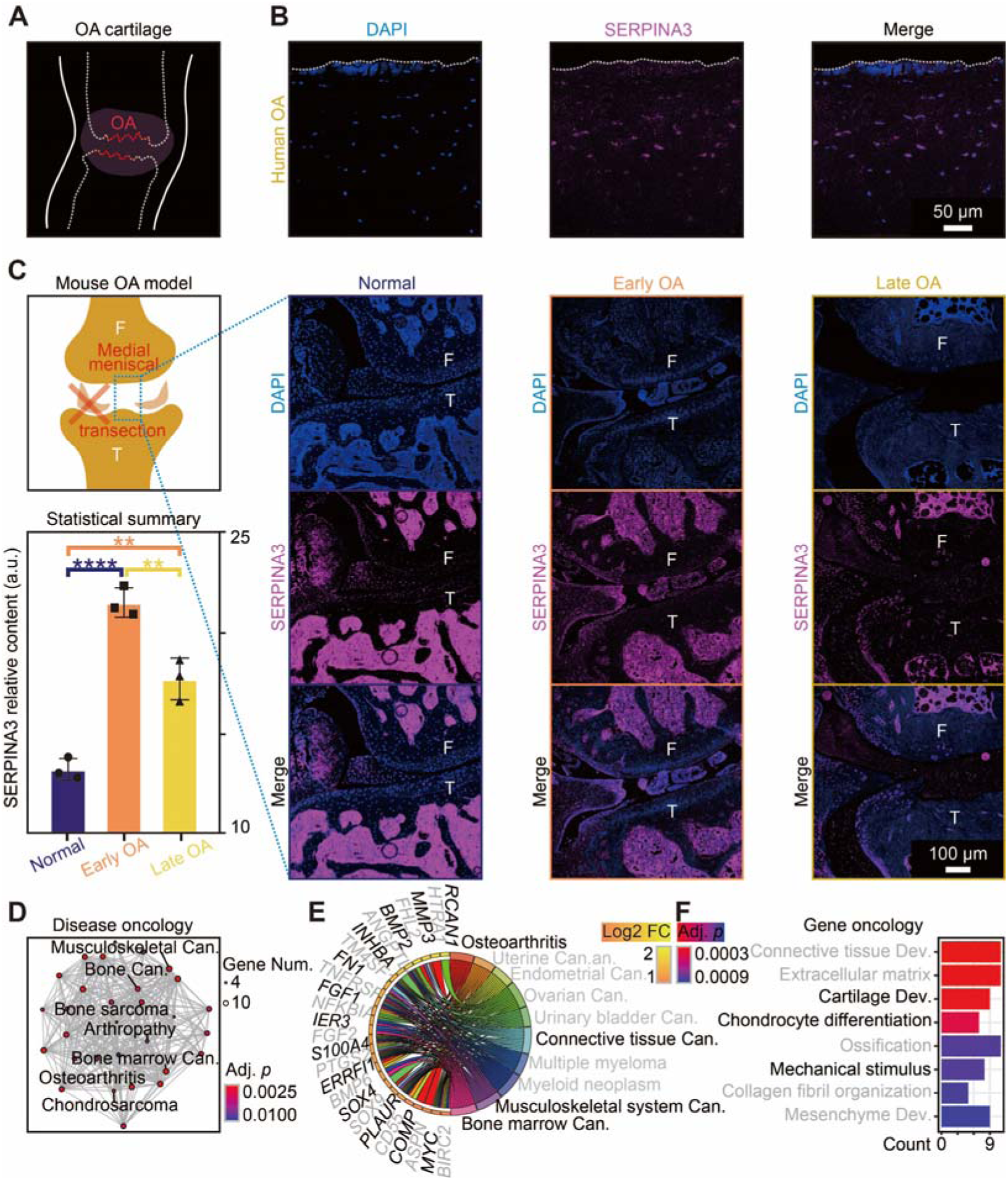
Arthritic transcriptional change in SERPINA3^+^ chondrocytes. (A) Schematic diagram showing degenerated articular cartilage surface in OA patients. (B-C) Anatomical sampling and immunostaining of DAPI (blue) and SERPINA3 (red) for chondrocyte subtype 14/15 in human OA samples (B) and mouse MMT-OA samples (C). On the left, schematic diagram showing the mouse MMT-OA model, and bar plots demonstrating the significantly elevated protein level of SERPINA3. Abbreviations including femur (F), tibia (T), and arbitrary unit (a. u.). (D-F) Enrichment analysis for DEGs in OA (Fig. S3A) through the DO (D-E) and GO (F) in chondrocyte subtype 14. Abbreviations including adjusted (Adj.), fold-change (FC), cancer (Can.), and development (Dev.).

The proliferation and differentiation of chondrocytes are critical for the treatment of OA joint degeneration. Recent studies have identified SERPINA3 as a marker of cartilage differentiation, playing a pivotal role in regulating the extracellular matrix (ECM) during fetal cartilage development [43]. When SERPINA3 is silenced, the chondrogenic markers (pAkt/SOX9/COLII/ACAN) significantly decrease, leading to reduced cartilage particle formation [43]. Additionally, SERPINA3 inhibits the enzymatic activity of arginyl collagenase by preventing proteolytic degradation, thereby halting cartilage matrix breakdown [41]. Collectively, these findings underscore the role of SERPINA3 in promoting chondrogenesis, regulating ECM dynamics and finally alleviate OA inflammation. However, the function of SERPINA3^+^ chondrocytes (chondrocyte subtype 14/15) in OA remains insufficiently investigated.

The existence of SERPINA3^+^ chondrocytes was validated in the articular cartilage of human late OA knee samples (Fig. 3B). To explore the involvement of SERPINA3^+^ chondrocytes in OA development, we established an OA mouse model via medial meniscal transection (MMT) surgery. Our results demonstrated a significant elevation of SERPINA3 in early OA (Fig. 3C). The observed retraction of SERPINA3 from early to late OA may be attributed to the severe cartilage loss in late OA, or possibly due to the exhaustion of SERPINA3^+^ chondrocytes. Further research is required to interpret their functions in OA development.

To elucidate the molecular networks, we examined the differentially expressed genes (DEGs) between OA and healthy controls (intact cartilage) in SERPINA3^+^ chondrocytes (Fig. S3A-D; Table S3). Disease ontology (DO) enrichments revealed significant associations with OA (adjusted *p* = 0.000), arthropathy (adjusted *p* = 0.006), bone marrow cancer (adjusted *p* = 0.000), and chondrosarcoma (adjusted *p* = 0.002) (Fig. 3D, E and Fig. S3E, F). DEGs involved in OA contained *ASPN*, *BMP2*, *CILP*, *COMP*, *CRYAB*, *ERRFI1*, *FHL2*, *FN1*, *HTRA1*, *HMOX1*, *HTRA1*, *MMP3*, *PTGS2*, *SERPINE1*, *SOX9,* and *TNFRSF11B*, which served as druggable targets for precision medicine (Table S3).

For further decoding of the intrinsic regulation, we conducted Kyoto encyclopedia of genes and genomes (KEGG) and gene ontology (GO) enrichment analysis (Table S3). The enrichment analysis revealed pathways significantly associated with cartilage development (adjusted *p* = 0.000), chondrocyte differentiation (adjusted *p* = 0.000), response to mechanical stimulus (adjusted *p* = 0.001), TNF signaling pathway (adjusted *p* = 0.020), Hippo signaling pathway (adjusted *p* = 0.043) (Fig. 3F and Fig. S3). These pathways have been reported with connection to chondrogenesis and OA development [44, 45]. In summary, we uncovered OA-elevated SERPINA3^+^ chondrocytes and provided new insights into the molecular mechanisms underlying OA pathogenesis.

### SERPINA3 promotes chondrogenesis in human OA chondrocytes

SERPINA3 has been identified as a key regulator of chondrogenesis and represents a promising therapeutic target for OA management [22, 34, 41–43]. In this study, to validate the chondrogenesis-promoting effect of SERPINA3, we took human pluripotent stem cell (hPSC)-derived chondrocytes as an in vitro model and simulated OA condition by adding IL-1β (Fig. 4A) [46, 47]. The hPSCs were sequentially differentiated into somite cells, sclerotome cells and chondral pellets. Then the chondral pellets were digested and plated for lentivirus transduction (Fig. 4A). Immunostaining of SOX9 and COLII revealed the expression of chondral markers in the induced chondrocytes, indicating the fidelity of the in vitro induction (Fig. 4B).

**Figure 4.**
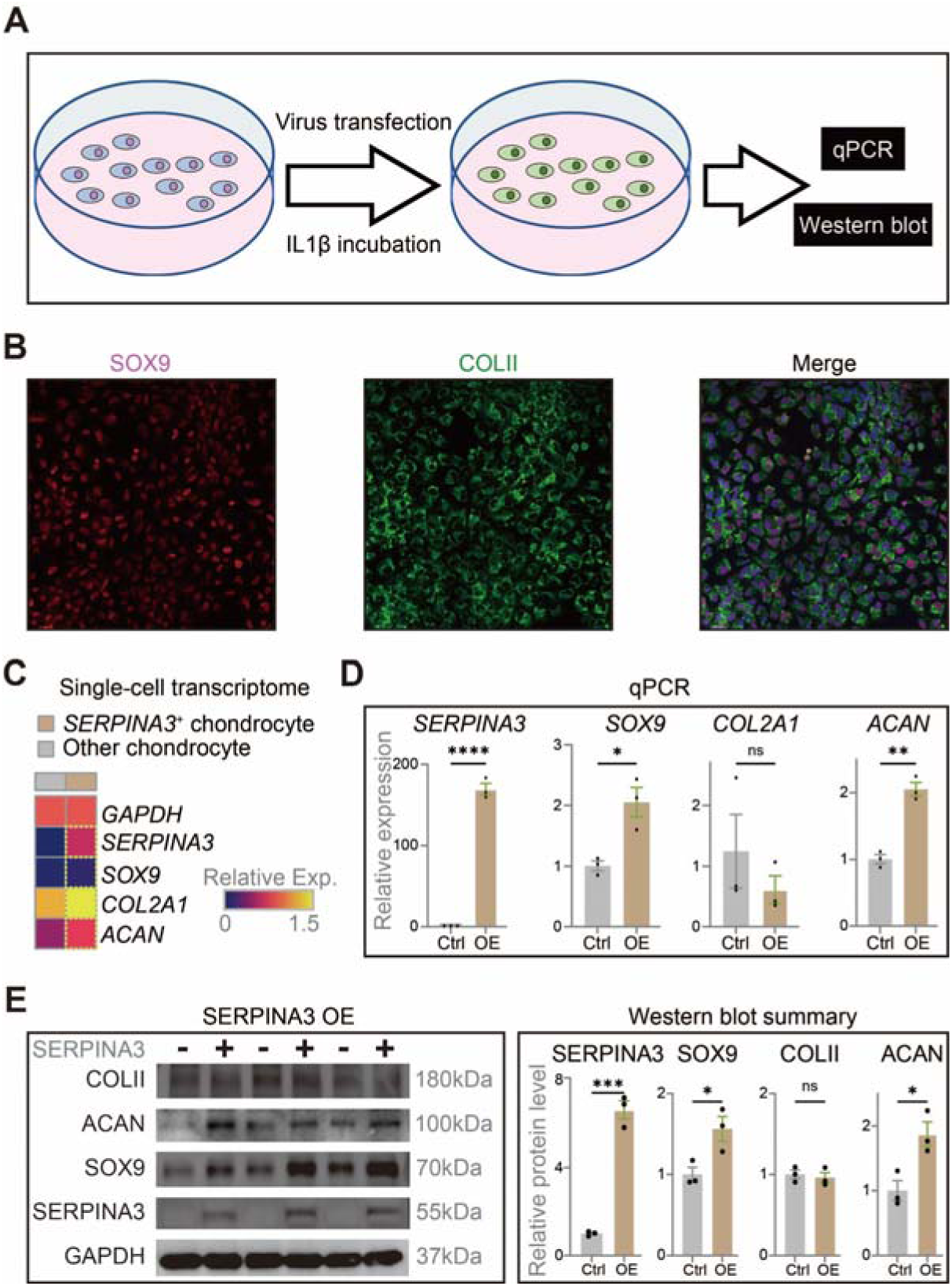
SERPINA3 promotes chondrogenesis in human OA chondrocytes. (A) Schematic diagram showing the in vitro experiments on human OA chondrocytes. (B) Immunostaining of SOX9 (red) and COLII (green) in hPSC-derived chondrocytes. (C) Heatmap showing the average expression normalized by the expression of GAPDH of *SERPINA3*^+^ chondrocytes and other chondrocytes under OA condition. (D) Bar plots showing the results of quantitative polymerase chain reaction (qPCR). (E) Left photographs and right bar plots showing the results of Western blot normalized to GAPDH expression. Abbreviation including kilodalton (kDa).

The SERPINA3^+^ chondrocytes were identified with a higher chondrogenesis pattern (*SOX9*, *COL2A1*, and *ACAN*) under OA condition (Fig. 4C) [47]. Thus, we supposed that SERPINA3 could promote the expression of chondrogenesis-associated and ECM-associated genes. Through virus transfection, we detected over 150-fold and 5-fold increases of SERPINA3 in transcriptional and protein levels (Fig. 4F, E and Fig. S4A, B). The significant upregulation of SOX9 (as chondrogenesis-associated transcription factors) and ACAN (as ECM-associated genes) matched reported findings (Fig. 4D, E and Fig. S4B-E). However, the transcript and protein level of COLII had no significant change, potentially due to the differences between primary chondrocytes and hPSC-derived chondrocytes (Fig. 4D, E and Fig. S4D). Collectively, SERPINA3 overexpression promoted chondrogenesis and served as a therapeutic target to alleviate OA [22, 34, 41–43].

### Microenvironment and extracellular matrix

As shown in Fig. 1A, JointMap categorized six major populations, including chondrocytes, progenitors, muscle cells, VSMCs, EndCs, and ImmCs. The intercellular communication within the joint microenvironment critically influences development, homeostasis, regeneration, aging, and pathological conditions [1, 2, 48]. Functional enrichment of chondrocyte subtype 14/15 (SERPINA3^+^ chondrocytes) identified cartilage development (adjusted *p* = 0.040), cell-cell signaling (adjusted *p* = 0.000), and extracellular space (adjusted *p* = 0.037) (Table S2). These molecular signatures collectively indicate substantial microenvironmental remodeling.

Intercellular interactions under diverse pathological conditions were systematically investigated to identify potential therapeutic targets. Crosstalk was decoded based on ligand-receptor pairs (Fig. 5A and Fig. S5A) [49]. Specifically, we examined expression difference of ligands and receptors in SERPINA3^+^ chondrocytes compared to other chondrocytes (Fig. 5B and Fig. S5B). Notably, receptor NCL exhibited significantly elevated expression in SERPINA3^+^ chondrocytes under OA conditions (Fig. 5B). This finding was further corroborated by immunofluorescence staining of human OA articular cartilage, which demonstrated the colocalization of SERPINA3 and NCL proteins (Fig. 5C). To validate these observations in vivo, we employed the MMT-OA mouse model to characterize microenvironmental alterations during OA progression. It unveiled significantly enhanced crosstalk (elevated NCL proteins) within SERPINA3^+^ chondrocytes (Fig. 5D and Fig. S5C). Together, these findings strongly suggest that therapeutic researches of the OA joint microenvironment should target these altered cellular interactions.

**Figure 5.**
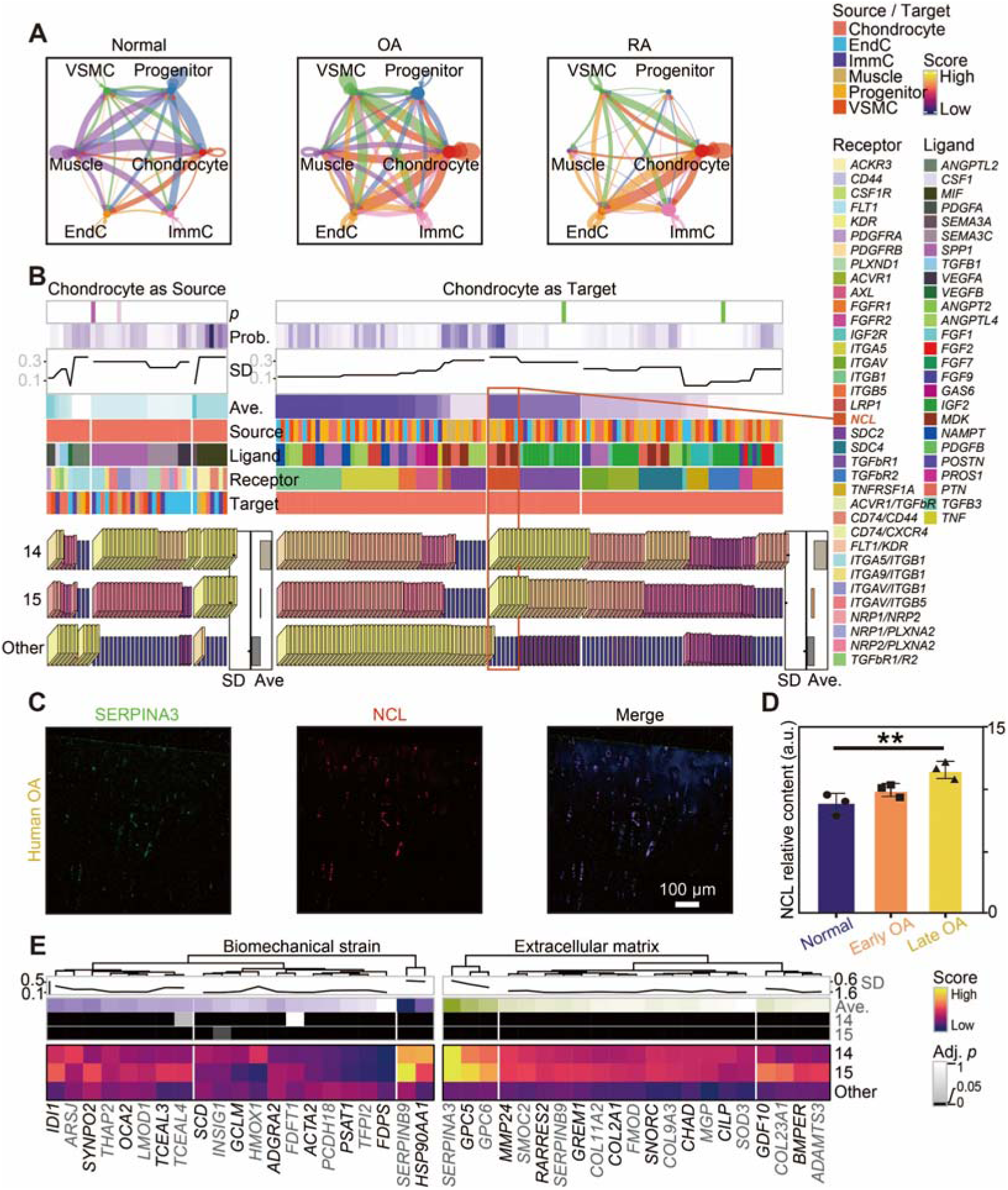
Microenvironment and extracellular matrix. (A) Circular dot plots presenting the crosstalk among cell types of the chondral microenvironment under normal, OA, and RA conditions speculated by CellChat [49]. The arrow from source cells (expressing ligands) pointing at the target cells (expressing receptors). The edge width representing the normalized numbers of ligand-receptor pairs. (B) Annotation heatmaps on the top showing detailed information, including *p* value, probability (Prob.) of cell–cell communication, standard deviation (SD), and average (Ave.) expression. Heatmaps presenting the scores (normalized average expression of chondrocytes) of genes (ligands in the left heatmap or receptors in the right heatmap). (C) Anatomical sampling and immunostaining of DAPI (blue), SERPINA3 (green) for chondrocyte subtype 14/15, and NCL (red) in human OA samples. (D) Bar plots demonstrating the elevated protein level of NCL in mouse MMT-OA model. (E) Heatmaps presenting the scores (normalized average expression of chondrocytes) of genes related to biomechanical stain (GSE165874; left) and ECM from AmiGO (right) [51].

Except cellular level, the impact of OA on joints could also be observed in ECM and biomechanical stress [5, 6]. Based on the pubic data under cyclic tensile treatment (GSE165874), we investigated mechanical-strain-responding genes in SERPINA3^+^ chondrocytes (Fig. 5E). Those significantly changed expression of biomechanical stress implicated the potential alteration of ECM. As a complex three-dimensional macromolecular network, ECM provides structural support for articular cartilage [1, 50]. Our results demonstrated substantial upregulation of ECM-related genes within SERPINA3^+^ chondrocytes (Fig. 5E and Fig. S5D). Collectively, these findings characterized the molecular alterations in the OA microenvironment and provided therapeutic targets.

### Multi-omic integration of chondrocytes in JointMap

OA is a globally prevalent chronic disabling disorder, affecting over 50% of individuals aged 65 or older [7]. Our study demonstrated that SERPINA3^+^ chondrocytes exhibited a more aged sample composition (Fig. S2E). This finding implied that SERPINA3^+^ chondrocytes possessed age-dependent accumulation (Fig. S2E). The impact of aging on chondrocytes needs further exploration. Transcriptional dynamics are crucial for gene expression regulation [52]. To address this gap, we performed kethoxal-assisted single-stranded DNA sequencing (KAS-seq) on chondrocytes under aging experiments. This approach provides direct reads of transcriptional activity with high temporal resolution [52, 53].

We created two models to simulate the effects of aging: first, we compared transcriptional activity between passage (P) 2 and 13 of rat primary chondrocytes isolated from femoral condyles (Fig. 6A); and second, we examined the inflammation induced by IL-1β (10/30/60 minutes) on chondrocytes in vitro (Fig. S6A) [54]. Pearson correlation analysis revealed significant correlations among inflammation groups (Fig. 6B). Visualization of KAS-seq data confirmed the aggregation of the “input” group that served as a blank control (Fig. 6C). Additionally, little influence of various cell lines was verified (Fig. S6B). Together, these results confirmed the quality of KAS-seq data.

**Figure 6.**
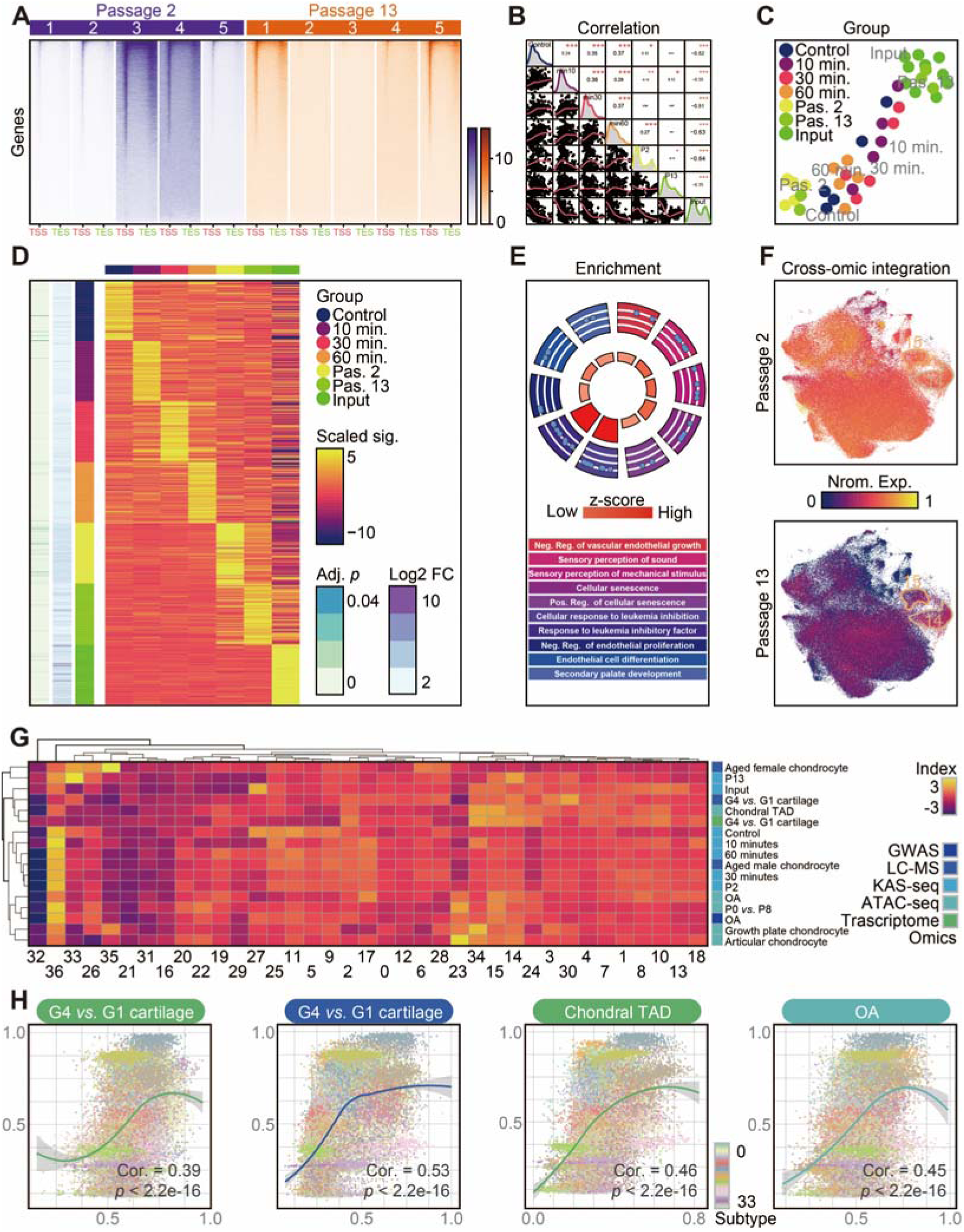
Multi-omic integration of chondrocytes in JointMap. (A) Heatmap showing KAS-seq signal distribution at gene-coding regions for P2 and P13 chondrocytes. Regions 3 kb upstream of transcription start site (TSS) and 3 kb downstream of transcription end site (TES) were shown. (B) Asterisk plots (up) and scatter plots (down) showing the correlation between samples of different experimental groups. (C) Embedding showing samples of different experimental groups. (D) Heatmaps presenting the top 100 differential signal (Sig.) of KAS-seq, with adjusted *p* values and log2 fold-change annotated on the left. (E) Dot plots of enrichment results based the genes shown in Fig. 6D of P13. Abbreviations including positive (Pos.), negative (Neg.), and regulation (Reg.). (F) Embedding showing normalized weighted expression of KAS-seq signal based the genes shown in Fig. 6D, with chondrocyte subtype 14/15 circulated. (G) Heatmap presenting the normalized average expression of multi-omic annotations in chondrocyte subtypes. (H) Dot plots presenting chondrocytes colored by subtypes. The correlation was calculated between OA scores (based on clinical diagnosis) and the integration of multi-omic data (annotated on the top).

To interpret genes with specific transcriptional activity (Fig. 6D), we applied GO enrichment analysis. Over-expansion (P13 compared to P2 chondrocytes) elevated pathways including sensory perception of mechanical stimulus and cellular senescence (Fig. 6E and Fig. S6C). Notably, the impact of inflammation (IL-1β) was captured quickly within the first test (10 minutes) and exhibited post-transcriptional gene silencing effects (Fig. S6D). Collectively, the impact of over expansion of chondrocytes in elderly individuals (P13 *vs*. P2) and the influence of inflammation in OA (simulated by IL-1β) were preliminarily unveiled by KAS-seq.

To integrate these omic data, we incorporated multi-omic datasets into the JointMap (Table S1), including liquid chromatography-mass spectrometry (LC-MS) [55, 56], assay for transposase-accessible chromatin with high-throughput sequencing (ATAC-seq) [57–60], and KAS-seq. For instance, we computed weighted indexes of KAS-seq for P13 based on log2 fold-change and gene symbols (Fig. 6F). The integration shown high accumulation of P13 indexes in chondrocyte subtype 14/15 (Fig. 6F). This indicated that the aged composition of SERPINA3^+^ chondrocytes may be associated with long-term continuous loss and regeneration of articular cartilage. However, interpreting cross-omic integration results is challenging [23]. We identified variations caused by subtypes with small cell numbers, such as chondrocyte subtype 32 and 36 (Fig. 6G). Interestingly, a high correlation (r = 0.83) was observed between “LC-MS aged male chondrocyte” indexes and “KAS-seq 60 minute” indexes (Fig. S6E). This is potentially arising from overlapping regulatory pathways involving transcriptional activity and proteins.

Furthermore, OA scores derived from clinical diagnosis exhibited significant correlation with multi-omic indexes, including “grade (G) 4 *vs*. G1 cartilage” indexes, “chondral topologically associating domain (TAD)” indexes, and “OA” indexes (Fig. 6H). These findings indicated the shared biological pathways in different omic levels. However, elucidating the genome-transcriptome-proteome regulatory networks remains a critical scientific problem. Our research provided a basic framework to integrate multi-omic data into JointMap.

### Spatial traits of chondrocyte subtypes

As mentioned, we could use “articular cartilage” or “hyaline cartilage” to terminologically name one chondrocyte. Similarly, the chondrocyte located in meniscus could be referred as “meniscal cartilage” or “fibrocartilage”. These overlapped classification standards sometimes result in confusion [2–4, 8–21]. To address this, we employed high-resolution gradient clustering in big data with machine learning algorithms to impel the establishment of a universal classification standard. Moreover, the homology and variations between meniscal and articular cartilage was analyzed based on our previous data [4, 40]. However, many studies focus exclusively on one isolated cartilage tissue. Together, the heterogeneity of chondral tissues has been widely recognized, yet comprehensive mappings for the distribution across various anatomical tissues was necessitated [1–3].

JointMap addresses this gap by compiling data from a wide range of chondral tissues, enabling the identification of spatially specific subtypes (Fig. S7A, B; Table S1). The compositional variability indicated subtype-specific localization preferences (Fig. 7A). For instance, chondrocyte subtype 1/8/10/13 were predominantly localized in both talus and articular cartilage (Fig. 7A). Notably, most subtypes exhibit multi-regional distribution patterns spanning three or more anatomical sites (Fig. 7A).

**Figure 7.**
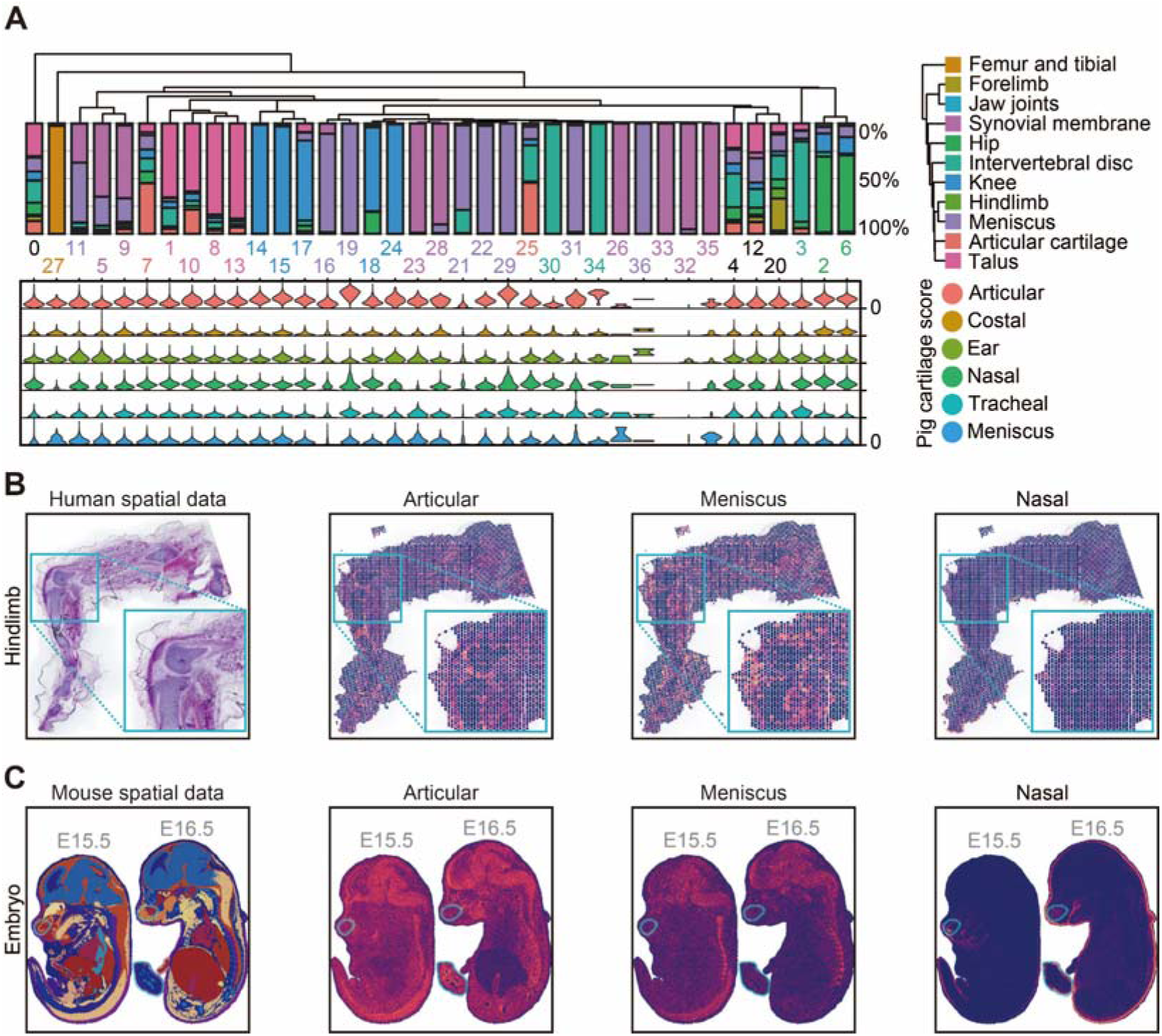
Spatial traits of chondrocyte subtypes. (A) Bar plots on the top presenting the compositional percentage of each chondrocyte subtype (corresponding to Fig. S7A). Bar color representing the source of the sequenced chondral tissues. Violin plots on the bottom showing the spatial module scores based on DEGs of pig cartilaginous tissues. Violin color representing the source of the specific location (corresponding to Fig. S7C). (B-C) Visualization of the spatial transcriptome of the 8.1 PCW human hindlimb sample (B) [2] and embryonic mice of the embryonic day 15.5 (E15.5) and E16.5 (C) [61] showing the spatial module scores of diverse pig cartilaginous tissues.

Additionally, we developed spatial module scores for different chondrocyte subtypes using our transcriptional data from various porcine cartilaginous tissues (Fig. 7A and Fig. S7C). For instance, chondrocyte subtype 19/29/31 exhibited elevated meniscal spatial module scores, consistent with exclusive composition of menisci. Conversely, chondrocyte subtype 11/16/21/22 showed low meniscal spatial module scores. This contrast may be caused by the heterogeneity of meniscal cartilage: our pig cartilaginous tissues lacked single-cell resolution, and meniscal cartilage may contain chondrocytes other than fibrocartilage (marked by COLI) [4, 40]. Overall, these quantitative scores provided insights into the subtype localization patterns.

Validation of these spatial module scores was performed using independent spatial transcriptomics datasets from human hindlimb tissues and murine embryonic specimens (Fig. 7B, C). Notably, articular spatial module scores demonstrated robust enrichment in human articular cartilage and murine footpads (Fig. 7B, C). Meniscal spatial module scores concentrated at the meniscal interface of articular cartilage in human hindlimbs (Fig. 7B), while nasal spatial module scores specifically marked the nasal region of embryonic mice (Fig. 7C). In conclusion, these cross-species validations confirmed the biological relevance and applicability of JointMap spatial scoring system.

In summary, cartilage is a heterogenous tissue and consisted of diverse subtypes. And one chondrocyte subtype could distribute across various anatomical tissues. Understanding the exact relation of subtypes and their locations requires further systematic study.

### Tissue-specific subtypes in JointMap

The remarkable cell compositional heterogeneity and spatial specificity unveiled by JointMap have motivated us to construct a comprehensive tissue-specific atlas. By analyzing articular cartilage and synovium [8], we noticed a blurring distinction between profiled synoviocytes (synovial fibroblasts) and chondrocytes. To further investigate this, we incorporated synovial membrane (synovial tissue) data into JointMap (Fig. 8A). Our analysis confirmed that synoviocytes and chondrocytes did not exhibit a clear separation, which implicated the absence of distinct molecular boundaries between these cell populations (Fig. 8A). We calculated synovial scores in single-cell level using synoviocyte-related markers identified in previous studies (Fig. S8A) [8, 15, 62, 63]. Cells with elevated synovial scores recapitulated the distribution characteristics of synoviocytes (Fig. 8A and Fig. S8A). Collectively, this metric served as a reliable discriminative criterion for synoviocyte identification.

**Figure 8.**
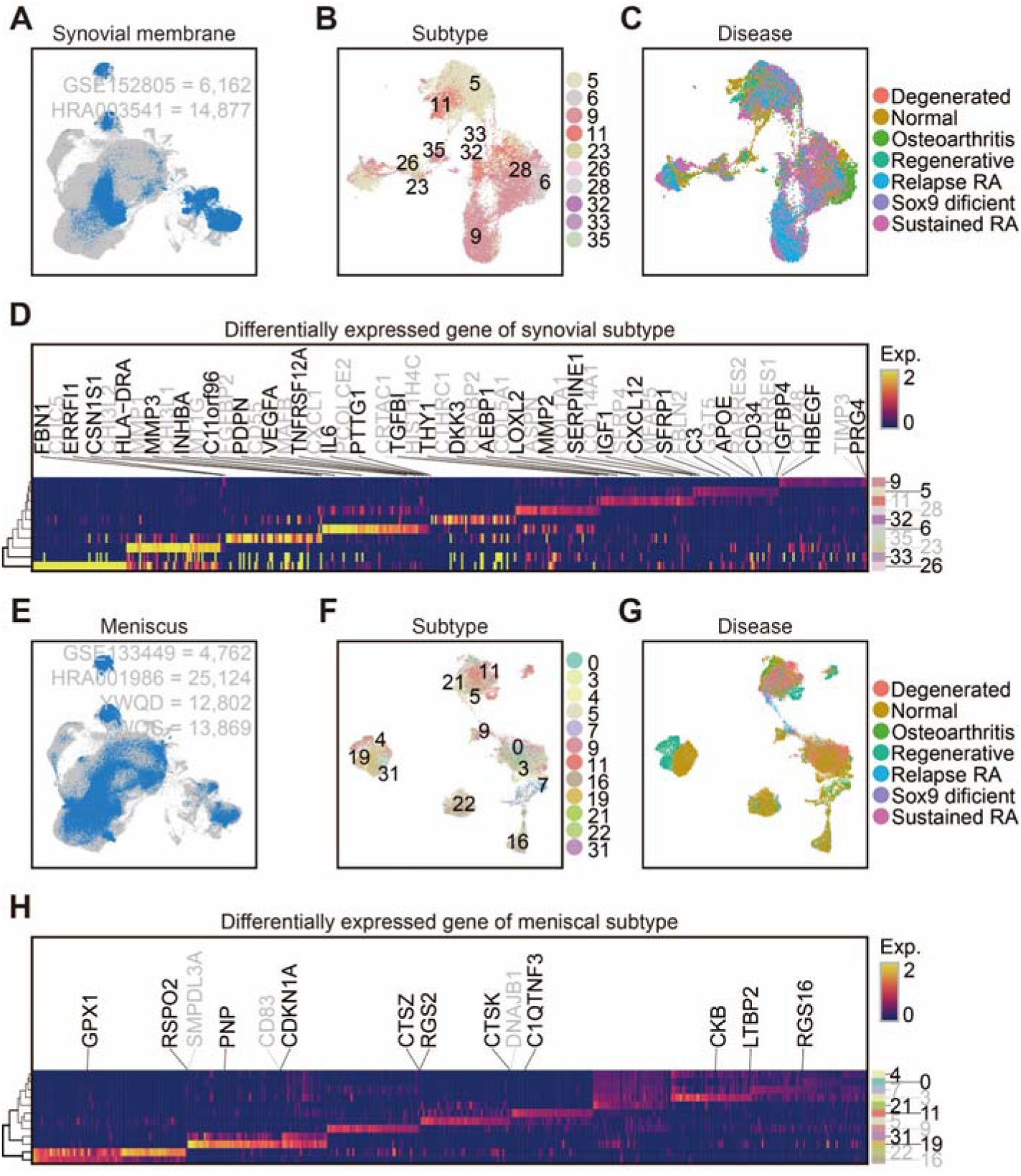
Exploration of synovial and meniscal cells. (A) Embedding showing all cells with these from synovial tissue colored in blue [8, 15]. (B) Embedding showing synoviocytes annotated by subtype identities. (C) Embedding showing synoviocytes annotated by the disease conditions of the sequenced samples. (D) Heatmap presenting the DEG expression of diverse synovial subtypes, with previously reported synovial-subtype-related genes annotated [8, 15, 62, 63]. (E) Embedding showing all cells with these from menisci colored in blue [4, 9, 12, 40]. (F) Embedding showing meniscal chondrocytes annotated by subtype identities. (G) Embedding showing meniscal chondrocytes annotated by the disease conditions of the sequenced samples. (H) Heatmap presenting the DEG expression of diverse meniscal subtypes, with meniscal specific genes annotated.

We systematically isolated synovial cells specifically from the synovial membrane or exhibiting elevated synovial scores (Fig. S8B) [8, 15]. The visualization demonstrated pronounced molecular heterogeneity (Fig. 8B). These synoviocyte subtypes exhibited differential enrichment in diseases (Fig. 8C and Fig. S8C). This observation aligned with previously reported synoviocyte subtype features (Fig. 8D). However, to connect the disease conditions to synovial transcriptional networks, more sophisticated datasets should be created.

Applying analogous methodologies to porcine cartilaginous tissues, we identified a panel of meniscal markers. Cells with high meniscal spatial module scores recapitulated the distribution characteristics of meniscal chondrocytes (Fig. 8E and Fig. S8D). We subsequently isolated meniscal chondrocytes gathered specifically from the menisci or exhibiting elevated meniscal spatial module scores (Fig. S8E) [4, 9, 12, 40]. Further characterization of meniscal chondrocyte subtypes with diseases required further investigation (Fig. 8F, G and Fig. S8F). DEG analysis identified subtype-specific transcriptional alterations associated with meniscal pathophysiology (Fig. 8H).

To decode the influence of diseases on the menisci, we employed the mouse MMT-OA model and the mouse meniscal regeneration model [4, 40]. Although quantitative proteomic analysis failed to detect significant change in CTSK (the marker of meniscal chondrocyte subtype 11), the results revealed a progressive increase from control to advanced disease states in mouse menisci (Fig. S8G). Complementing these findings, our regeneration model demonstrated preferential enrichment of CTSK^+^ chondrocytes in the regenerated zones of menisci (Fig. S8H). Collectively, these findings suggested a critical role of CTSK^+^ chondrocytes in OA and regeneration.

## DISCUSSION

JointMap, a unified transcriptomic reference atlas focusing on chondral tissues, integrated 545,946 cells and 22,161 genes. It served as a crucial resource for dissecting druggable networks targeting articular diseases. The cellular heterogeneity of JointMap was validated through immunostaining in both human samples and animal models and in vitro OE cell culture. We identified OA-elevated chondrocyte subtype 14/15 (SERPINA3^+^ chondrocytes) with upregulated intercellular communication, which unveiled potential therapeutic targets for clinical studies.

To construct JointMap, 14 chondral datasets were selected and integrated using the widely recognized algorithms, Harmony and Seurat. Additionally, our study employed reciprocal principal component analysis (PCA), joint PCA, canonical correlation analysis, and reciprocal latent semantic indexing, all of which yielded positive results [21, 25]. Despite the variations in sequencing strategies over time (Smart-seq *vs.* 10× Genomics Chromium; single-cell *vs.* single-nucleus), we implemented rank-based normalization for each sample to minimize batch effects in the count matrix [25, 64]. We encouraged future studies to develop informatic methods (e.g., artificial intelligence) to minimize the batch effect and explore key molecular networks. Overall, JointMap provides a unified framework for the systematic annotation of chondral tissues across multiple species.

The collected animal datasets offered valuable insights into interspecies conservation and evolution. In this study, we mainly consolidated data from human samples, along with data from dog, sheep, mouse, and zebrafish. Here, we decoded the interspecies conservation and evolution through gene regulatory networks. However, only these subtypes with sufficient number of cells were selected. Future research could expand JointMap to include more species, ultimately facilitating the creation of comprehensive cross-species chondral data.

The adult human samples were typically collected under severe disease conditions (*e*.*g*., synovial joint samples from total knee arthroplasty in late-stage OA patients). The young and healthy control samples are of extreme deficiency. In JointMap, the control samples majorly were collected from intact lateral condyle of HRA002569 (Table S1) [13]. And we calculated disease-specific scores (e.g., OA scores) using the clinical diagnosis. However, a high OA score of one cell only suggested its potential relation to OA. To further support this, we conducted multi-omic analysis and validated that OA scores significantly correlated with “G4 *vs*. G1 cartilage” indexes, “chondral TAD” indexes, and “OA” indexes. These scores and indexes implicated shared common biological pathways across multiple omic levels.

Datasets from diverse cartilage tissues can enhance our understanding of chondrocyte heterogeneity. In this study, we developed spatial module scores to link chondrocyte subtypes with their potential locations. Furthermore, we developed two location-specific maps for synovial and meniscal research. Most datasets in JointMap pertained to synovial joint (knee), and more chondral tissues are needed (*e*.*g*., shoulder joint [3]).

SERPINA3, also known as alpha-1-antichymotrypsin, is a secreted protein. It primarily inhibits serine protease by preventing proteolytic activity, which results in ECM changes [41]. As an acute inflammatory response protein, the plasma SERPINA3 protein concentration correlated with the severity and duration of inflammation [41, 42]. Its expression is consistent with the inflammatory response observed in OA. *SERPINA3* is widely expressed in various tissues, including the in various tissues, including the brain, liver, kidney, gallbladder, pancreas, and other tissues [41, 65]. And its upregulation has been noted in heart failure, nervous system diseases and cancer [41, 65]. Additionally, the gender difference in its expression (higher in females than in males) resembles the incidence of OA (more common in women than in men). And the recent publication using genome-wide association study on 1,962,069 individuals has identified SERPINA3 as one of effector genes involved in OA [34].

At the genomic level, human (*SERPINA3*) and mouse (*Serpina3n*) share 61% homology [35, 36]. We found that SERPINA3^+^ chondrocytes were not specific to humans. More importantly, we validated the expansion of these subtypes in human patient and mouse OA-models via immunostaining. Recently, SERPINA3 has been reported to functions in neocortical folding and cognitive improving during evolution [35, 36]. In the osteochondral system, SERPINA3 is a marker of cartilage differentiation [43]. It may be an upstream of SOX9, which is important for cartilage formation [43]. The physical interaction at the protein level is 77.64% for human (SERPINA3-SOX9) and 45.00% for mouse (Serpina3n-Sox9) [66]. And the predicted functional partners for SERPINA3 include KLK3, CTSG, CTRB2, CTRB1, and A2M [67]. Furthermore, the “blood microparticle” (GO:0072562) pathway is enriched in protein networks, machine-learning features and OA-elevated genes of chondrocyte subtype 14 [67].

Recent studies have identified SERPINA3 as a marker of cartilage differentiation and a critical regulator of ECM-related genes during early chondrogenesis. In the osteochondral system, *SERPINA3* is primarily expressed during embryonic cartilage development [43]. In OA treatment studies, upregulation of SERPINA3 via Sinensetin exhibits reduction of inflammation and cartilage damage [68]. Our SERPINA3 OE in OA chondrocytes validates the upregulation of chondrogenesis. Combining our findings, we propose that SERPINA3^+^ chondrocytes have a positive impact on chondrocyte differentiation and OA treatment.

## MATERIALS AND METHODS

### Single-cell RNA-seq data processing

Single-cell data were downloaded (Table S1) [2, 8–17] and processed following our previous research [4, 25, 40, 69]. The sequenced data was processed using ‘CellRanger v7.1.0’ package in ‘Python v3.10.15’ with the reference genome *Homo sapiens* (human; GRCh38) or *Mus musculus* (mouse; mm10) to generate filtered expression matrices which were then analyzed with ‘Seurat v5.0.1’ package in ‘R v4.2.3’ [23]. The sample with the cell number higher than 1,000 was kept for downstream analysis. Except human data, all genes that can be transformed to human homologs (one to one) via ‘getLDS’ function from ‘biomaRt v2.54.1’ package and presented in at least 8 datasets were adopted. After normalizing the data, we scaled genes presented in at least 10 studies and in the chosen 3,500 highly variable features (via ‘FindVariableFeatures’ function from ‘Seurat’ package) [23].

The Harmony embedding based on 50 PCs were employed for UMAP visualization on purpose of eliminating batch effect (via ‘RunPCA’ and ‘RunUMAP’ functions from ‘Seurat’ package and ‘RunHarmony’ function from ‘Harmony v1.0.3’ package) [20, 21, 24]. Furthermore, reciprocal PCA, joint PCA, canonical correlation analysis and reciprocal latent semantic indexing were assessed, yielding positive outcomes [21, 25].

To further mitigate the batch effect in JointMap for these algorithms utilizing the count matrix, we conducted rank-based normalization for each sample across studies [25, 64]. In brief, single-cell level density scores of each major cell types (chondrocytes, progenitors, muscle cells, VSMCs, EndCs, and ImmCs) was calculated based on UMAP reduction via ‘kde2d’ function from ‘MASS v7.3-58.2’ package. Next, the cells with density scores higher than the mean were kept for downstream analysis. The genes of each sample were ranked and normalized to remove the unrepresentative genes with rank below the lower quartile. All the processed matrixes were merged together for these algorithms utilizing the count matrix [25, 64].

### Quality control

Following the initial data processing, cells with high mitochondrial percentage (more than 10) and low transcripts captured (count/cell less than 1,500 or feature/cell less than 500) were filtered out [25, 69]. Subsequently, we regressed out the effects of cell cycling to mitigate the influence of the G2M and S phases on data reduction and cell type classification. In detail, we used ‘CellCycleScoring’ and ‘ScaleData’ functions from ‘Seurat’ package to mitigate the effects of cell cycle heterogeneity by calculating cell cycle phase scores based on canonical markers to regress these out of the data [23].

Regarding contamination and doublets in scRNA-seq data, three packages were used to retain high-quality cells. Firstly, usage of droplet-based microfluidic devices would cause ambient RNA present in the cell suspension aberrantly counted, and would cause cross-contamination of transcripts between different cell populations. We performed the “decontX” function of ‘DecontX v0.99.3’ package to estimate and remove contamination in individual cells via Bayesian method [70]. Second, doublet data was created by the multiple cells captured within the same droplet or reaction volume. In detail, we used both the “computeDoubletDensity” function from ‘scDblFinder v1.12.0’ package [29] to identify potential doublet cells based on the local density of simulated doublet expression profiles and “doubletFinder” function from ‘DoubletFinder v2.0.4’ package [28] that generated artificial doublets from existing scRNA-seq data and defined which real cells preferentially co-localize with artificial doublets in gene expression space.

### Cell type classification

The major cell type identities were classified based on canonical marker genes [2, 4, 8–17, 25, 69]. Cells were further clustered based on the top 50 Harmony embeddings via ‘FindNeighbors’ and ‘FindClusters’ functions from ‘Seurat’ package with gradient resolutions [23]. To infer the molecular characteristics for each subtype, the l_2_-norm regularized logistic regression-based machine learning algorithm was applied to the rank-based normalized data via ‘LogisticRegressionCV’ function from ‘scikit-learn v1.3.2’ with modifications [31, 32]. To further investigate the transcriptional regulatory network of these cells, we calculated the regulon activity of transcription factors speculated via ‘grn’, ‘ctx’, and ‘aucell’ functions from ‘pySCENIC v0.12.1’ package [33].

The single-cell level disease-specific scores, including degenerated score, OA score, regenerative score, relapsed RA score, and sustained RA score, were calculated using the original metadata from the samples via ‘imbalance_score’ function from ‘bioc2020trajectories v0.0.0.93’ package (https://github.com/kstreet13/bioc2020trajectories).

### Functional analysis

The DEGs were extracted by ‘FindAllMarkers’ function from ‘Seurat’ package. To decode the intrinsic network of provided gene lists, DO, GO, and KEGG enrichment analyses were conducted via ‘enrichDO’ function from ‘DOSE v3.24.2’ package and ‘enrichGO’ and ‘enrichKEGG’ functions from ‘clusterProfiler v4.6.2’ package [71]. Additionally, the gene set enrichment analysis (GSEA) based on whole transcriptome was performed by ‘gseGO’ and ‘gseKEGG’ functions from ‘clusterProfiler’ package [71].

The crosstalk among diverse cell types in the chondral microenvironment under various conditions was examined using rank-based normalized data by ‘computeCommunProb’ and ‘aggregateNet’ functions from ‘CellChat v2.1.2’ package [49]. The enhanced ligand-receptor interactions were visualized via ‘Heatmap’ function from ‘ComplexHeatmap v2.14.0’ package [72].

The DEGs in response to mechanical strain through cyclic tensile strain treatment from public data (GSE165874) with adjusted *p* value lower than 0.05 and average log2 fold-change higher than 0.3 were calculated and used for visualization of OA-related subpopulations via ‘Seurat’ [23] and ‘ComplexHeatmap’ packages [72]. Furthermore, ECM-related genes were collected from AmiGO (https://amigo.geneontology.org/amigo/term/GO:0031012) [51].

### SERPINA3 promotes chondrogenesis in human OA chondrocytes

The human OA chondrocytes were induced as described previously with modifications [46, 47]. The H9 human embryonic stem cell line (WiCell, #WA09) was cultured in mTeSR1 medium (STEMCELL Technologies, #85850) on Matrigel (Corning, #354277). Cells were passaged every 3 days using EDTA (Procell, #PB180320) at a ratio of 1:6 and seeded with 5 μM Y27632 (Topscience, #T1870) for the first 24 hours. Cells were routinely tested to confirm there was no mycoplasma contamination. Differentiation of hPSCs into induced chondrocyte pellets was performed as described previously with modifications [46, 47]. Cells were dissociated into single cells using TrypLE Express (ThermoFisher, #12604021) and seeded at a density of 25,000 cells/cm^2^ on a Matrigel-coated plate. Basal differentiation medium (BDM) was prepared by adding 1× ITS-G (Procell, #PB180429) and 1× penicillin-streptomycin (Procell, #PB180120) into DMEM/F12 medium (Corning, #10-092CVRC). After 24 hours, differentiation was initiated using BDM supplemented with 3 μM CHIR99021 (Topscience, #T2310) for the first 48 hours. Medium was changed daily. For the next 48 hours, medium was switched to BDM supplemented with 200 nM LDN193189 (Topscience, #T6158) and 10 μM SB431542 (Topscience, #T1726). Medium was changed daily. Then the cells in one well of a 6-well plate were dissociated using EDTA and seeded in two Matrigel-coated 10-cm plate in BDM supplemented with 20 ng/mL FGF2 (SinoBiological, #10014-HNAE) and 300 nM SAG (Topscience, #T1779) for the next 48 hours to generate sclerotome cells. Medium was changed daily. The sclerotome cells were dissociated into single cells using TrypLE Express, and 500,000 cells in 1 mL chondrogenic medium (ScienCell, #7551) supplemented with 10 ng/mL TGF-β3 (SinoBiological, #10434-H01H) were pelleted at 300 g for 5 minutes in a 15-mL centrifuge tube. Pellets were cultured in the same medium for 21 days. Cells were fed with fresh medium every other day.

The OE of SERPINA3 in human OA chondrocytes was performed as described previously with modifications [47]. The generated induced chondrocyte pellets were dissociated into single cells using 0.2% collagenase II and seeded in Matrigel-coated 6-well plate at a density of 500,000 cells/cm^2^ with MEM-α medium (Cytiva, #SH30265.01) supplemented with 10% FBS (Corning, #35-078-CV). When the confluency reached about 60%, the cells were transduced with Lenti-SERPINA3-GFP or control Lenti-GFP lentivirus at multiplicity of infection about 10 with 4 μg/mL polybrene (Beyotime, #C0351-1mL) to ensure stable expression of SERPINA3. After 24 hours, the cells were washed with PBS and cultured in MEM-α medium supplemented with 10% FBS without lentivirus for 48 hours. Then the cells were treated with 10 ng/mL IL-1β (SinoBiological, #10139-HNAE) for 48 hours and harvested for further analysis.

### KAS-seq of chondrocytes under aging experiments

The process of KAS-seq and analysis were performed according to the modified manuscript [52]. In details, for rat primary chondrocytes isolation, the cartilage fragments were obtained from femoral condyles. The cartilage fragments were minced and digested with 0.1% type II collagenase (Invitrogen) at 37 °C for 12-16 hours with continuous shaking. Chondrocytes were resuspended in α Minimum Essential Medium (αMEM; Gibco) with 10% fetal bovine serum (FBS; Gibco, USA) and 1% penicillin/streptomycin (P/S; 100 units/mL penicillin and 100μg/mL streptomycin, HyClone). The primary chondrocytes were cultured in αMEM supplemented with 10% FBS and 1% P/S.

For replication-induced senescence, cells were passaged until they lost the ability of proliferation and became senescent. Chondrocytes were used as senescent cells after 15 passages. For inflammatory OA model, rat chondrocytes were starved for 4-6 hours without FBS and induced by IL-1β (10 ng/mL; 211-11B; PeproTech, Beijing, China) for 10, 30, and 60 minutes. Rat chondrocytes were labeled with N3-kethoxal using the same labeling protocol as described before. Briefly, chondrocytes were washed 3 times with PBS and treated with N3-kethoxal for 10 minutes. gDNA were isolated from chondrocytes by using PureLinkTM Genomic DNA Mini Kit (K182001; Thermo Fisher Scientific, USA) according to the manufacturer’s protocol.

After labelling, gDNA was fragmented using Tn5 transposase (TD501; Vazyme, China) at 37 °C, 500 rpm for 30 minutes. After biotinylation, gDNA was then purified using DNA Clean & Concentrator-5 kit (D4013; Zymo, USA) and 5 µL labelled DNA was retained as input. For bead-based enrichment, 5 µL of Dynabeads™ MyOne™ Streptavidin C1 (65,001; Thermo Fisher Scientific, USA) were washed three times using Binding and Wash buffer. The beads were mixed with 50 µL of the fragmented DNA obtained in the previous steps and slowly rotated at room temperature for 15 minutes. After incubation, the beads were washed five times with Binding and Wash buffer on a magnetic rack to remove the supernatant. The DNA conjugated beads and their respective inputs were used for library PCR. The library PCR was performed using i5 and i7 index primers (20027213; Illumina, USA) and NEBNext Ultra II Q5 Master Mix (M0544S; NEB, USA). The PCR reactions were heated for 5 minutes at 72 °C, followed by 10 minutes at 95 °C. The amplification was then performed for 15 cycles (10 seconds at 98 °C, 30 seconds at 60 °C, 1 minutes at 72 °C). The libraries were cleaned up by using MinElute PCR purification kit (28804; Qiagen, Germany). The data processing was based on the online codes and guidance [52].

### Cross-omic data integration

The analyzed tables containing the gene symbols and fold-change values were downloaded from the corresponding papers (Table S1). The indexes of the cross-omic data were calculated based on the weighted average expression in single-cell level of JoinMap. In details, “ATACseq_OA” indexes were created based on the ATAC-seq robust peak information from the knee joints of patients with osteoarthritis (the “table S3” in the original paper [57]). “ATACseq_Growth_plate_chondrocyte” and “ATACseq_Articular_chondrocyte” indexes were created based on the targeted motif testing growth plate/articular chondrocyte differentially accessible peaks from the knee joints of patients with osteoarthritis and mouse tissues (the “supplementary file 3f” in the original paper [58]). “ATACseq_Chondral_topologically_associating_domain” indexes were created based on the list of chondrogenic genes located in chondral topologically associating domains from the fetal chondrocytes isolated from mice bearing a Col2a1 fluorescent regulatory sensor (the “supplementary data 8” in the original paper [59]). “ATACseq_P0_vs_P8” indexes were created based on the chromatin landscape modifications in late-dedifferentiated (P0 *vs*. P8) chondrocytes from the primary hyaline chondrocytes isolated from uncalcified cartilage in knee joints of newborn mice (the “GSE193743_processed_data-4.csv.gz” in the original paper [60]). Samples from the severely damaged region were classified into the G4 cartilage. “LCMS_G4_vs_G1_cartilage” and “Trascriptome_G4_vs_G1_cartilage” indexes were created based on the DEGs and relative protein quantification of G4 cartilage comparing to G1 (the “supplementary table 1 and 3” in the original paper [55]). “LCMS_Aged_Female_Chondrocyte” and “LCMS_Aged_Male_Chondrocyte” indexes were created based on the abundance comparation between aged and young samples from the knee cartilage from male and female young, middle-aged, and aged C57/BL6 mice (the “Gabby_OTE_TMT16-frac-mouse-knee_20201117_proteins.xlsx” in the original paper [56]). “GWAS_OA” indexes were created based on the OA effector genes and their corresponding scores from the GWAS analysis based on 1,962,069 individuals (the “Supplementary Table 29: Effector genes and membership of highlighted pathways” in the original paper [34]).

### Validation in human samples and mouse OA model

The human and animal studies were approved by Peking University ethics committee (LA2021007). All applicable institutional and/or national guidelines for the care and use of animals were followed. In the present study, the human knee samples were harvested during total knee replacement of OA patients (female, 50-60 years old, race: Han nationality). The removed cartilage tissues were collected and fixed with 4% paraformaldehyde. The paraffin sections of 5 µm thickness were prepared for subsequent immunostaining.

A total of six mice (male, 2 months) was included. The total meniscectomy of medial meniscus was performed in one side of knee joint. The contralateral knee joints were used as normal samples. Briefly, the knee joint was exposed through regular medial parapatellar approach. The medial meniscus was removed using scissors. Three mice were executed at 4 weeks after operation. The samples were considered as early OA. Another three mice were executed at 12 weeks after operation. The samples were considered as late OA. The samples were fixed with 4% paraformaldehyde and decalcified using EDTA. The samples were embedded with paraffin. The 3 µm-thick paraffin sections were prepared using microtome (Leica, Germany). The sections were immersed into xylene and graded ethanol to deparaffinize and regain water. The heat induced antigen retrieval was completed using pH 6.0 citric acid for 20 minutes. The nonspecific protein binding was blocked using goat serum (Boster, AR0009, China) for 1 hour at room temperature. The sections were incubated with corresponding primary antibodies for 2 hours at room temperature. After thorough washing with PBST, the sections were incubated with corresponding secondary antibodies for 1 hour at room temperature, followed by DAPI incubation. Finally, the slices were sealed with anti-fluorescence quenching agent. The confocal microscope (Leica, Germany) was used to capture images. For semiquantitative analysis, the region of interest was drawn in the superficial cartilage zone of femoral condyle and tibial plateau. The integrated intensity of corresponding target was evaluated by Image J software (US National Institutes of Health, USA).

Antibodies utilized in this paper included SERPINA3 (Signalway antibody, 32122), CD31 (Proteintech 11265-1-AP), CD68 (Abcam ab303565), COL II (Abcam ab34712), CTSK (Abcam ab187647), and NCL (Active Motif 39541). The data were demonstrated with mean values ± standard deviation. The Shapiro-Wilk test was used to evaluate data distribution. The equal variance of data was checked before analysis. The ordinary one-way ANOVA with Bonferroni multiple comparison test was performed. The statistical significance was considered when the *p* value < 0.05. The statistical analysis was performed using ‘GraphPad Prism v8.0.1’ software.

### Spatial analysis

Mouse embryonic spatial data was downloaded and processed using ‘Seurat’ package [23, 61]. Human hindlimb spatial data was downloaded and processed via ‘pl.spatial’ function from ‘Scanpy v1.10.3’ package [2, 23, 73, 74].

The spatial scores (depicted in Fig. 7A and Fig. S8D) were calculated based on ‘cor’ function from ‘stats v4.2.3’ package and weighted average expression of DEGs from our numerous pig cartilaginous tissues. The single-cell level synovial scores were calculated based on genes associated with synovial subtypes identified in previous studies (*COL1A1*, *PDPN*, *THY1*, *HLA-DRA*, *IL6*, *CD34*, *CD55*, *DKK3*, *PRG4*, *TIMP3*, *CSN1S1*, *IGFBP4*, *CHI3L2*, *MMP2*, *MMP3*, *MMP1*, *MT1G*, *CXCL1*, *HBEGF*, *C11orf96*, *CRTAC1*, *ERRFI1*, *CLIC5*, *APOE*, *RARRES2*, *CHI3L1*, *COL14A1*, *GGT5*, *C3*, *SFRP1*, *RARRES1*, *MFAP5*, *FGFBP2*, *FBLN2*, *FBN1*, *ASPN*, *CRABP2*, *CD248*, *SFRP4*, *PCOLCE2*, *VEGFA*, *TGFBI*, *SERPINE1*, *CTHRC1*, *LOXL2*, *COL5A1*, *TNFRSF12A*, *AEBP1*, *MAFB*, *HSPA66*, *HIST1H4C*, *PTTG1*, *CENPF*, *CSN151*, *INHBA*, *WISP2*, *IGF1*, and *CXCL12*) via ‘AddModuleScore’ function of ‘Seurat’ package [8, 15, 23, 62, 63].

## CONCLUSIONS

In summary, we have systematically constructed JointMap, a practical reference atlas designed to explore druggable molecular networks for the treatment of articular disorders. We further performed KAS-seq and annotated other multi-omic data to elucidate the transcriptome of JointMap. Notably, our analysis validated the upregulated chondrogenesis function of SERPINA3^+^ chondrocytes under OA condition, highlighting these cells as promising candidates for future therapeutic targeting.

## MATERIALS AND METHODS

Detailed materials and methods are available in the Supplementary data.

## DATA AVAILABILITY

The JointMap data reported in this paper have been deposited in the OMIX, China National Center for Bioinformation/Beijing Institute of Genomics, Chinese Academy of Sciences (https://ngdc.cncb.ac.cn/omix: accession no.OMIX007049; https://ngdc.cncb.ac.cn/bioproject/browse/PRJCA028746) [75, 76]. The original datasets (except these from our laboratory) are available in the BioStudies database (http://www.ebi.ac.uk/biostudies) under accession number E-MTAB-8813, in the GSA-Human database (https://ngdc.cncb.ac.cn) under accession number PRJCA013514, PRJCA010181 and PRJCA008120, and in the NCBI-GEO database (https://www.ncbi.nlm.nih.gov/geo/) under accession number GSE160756, GSE216578, GSE184403, GSE169454, GSE162033, GSE152805, and GSE133449 [2, 8–17, 75–78].

## ACKNOWLEDGEMENTS

We would like to acknowledge the contributions and support of all staff of Center of Basic Medical Research, Institute of Medical Innovation and Research, Peking University Third Hospital for their assistance during the whole process of conducting this project. All national and institutional guidelines for the care and use of patient samples and laboratory animals were followed.

## AUTHOR CONTRIBUTIONS

H.W., W.Y., M.S. and Y.F. contributed equally to this work. H.W. collected and analyzed the bioinformatic data. W.Y. conducted the majority of the experiments. W.Y. and M.S. performed the in vitro culture of human OA chondrocytes. Y.F., L.C., and J.L. conducted KAS-seq. H.W., W.Y., M.S., Y.F., and X.H. prepared the manuscript. S.J., J.W., B.X., B.L., Y.D., J.C., and Y.A. conducted the surgery and sample collection.

## ADDITIONAL INFORMATION

**Competing interests:** The authors declare no competing interests.

**Ethical approval:** The human and animal studies were approved by Peking University ethics committee (LA2021007). All applicable institutional and/or national guidelines for the care and use of animals were followed.

---

**Figure S1.**
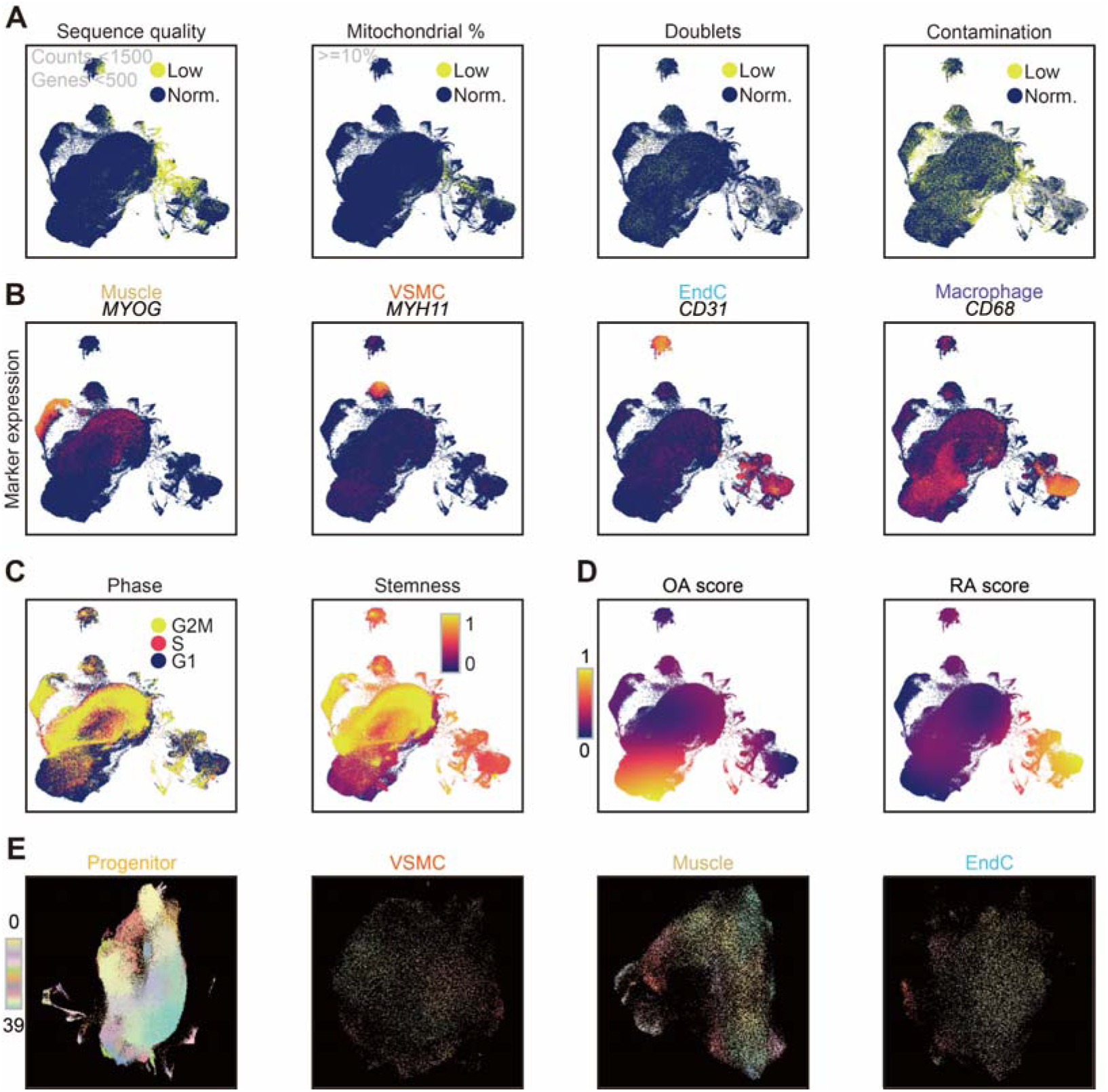
Unified chondral atlas JointMap. (A) Embedding showing sequence quality (cells with read counts less than 1,500 or gene number less than 500 marked as low quality), mitochondrial transcription percentage (cells with 10% or more as low quality), doublets (judged by both DoubletFinder [28] and scDblFinder [29]), and contamination (judged by DecontX [30]). Abbreviations including normal (Norm.) and low quality (Low). (B) Embedding showing expression of key markers. (C) Embedding showing cell cycle phases inferred by Seurat [23] and stemness scores inferred by CytoTRACE [26]. (D) Embedding showing simulated OA and RA scores. (E) Re-embedding of subtypes of progenitors, VSMCs, muscle cells, and EndCs.

**Figure S2.**
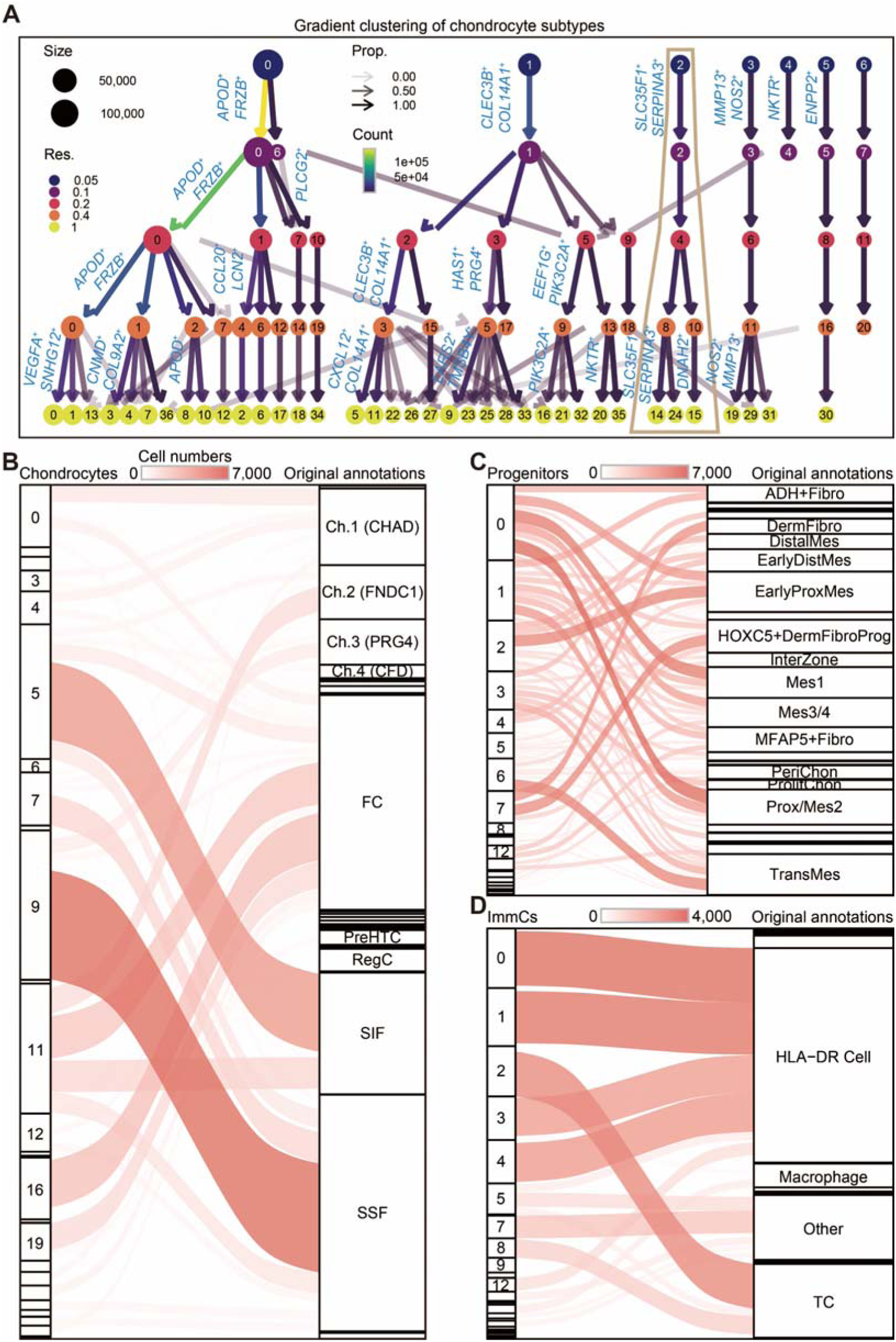
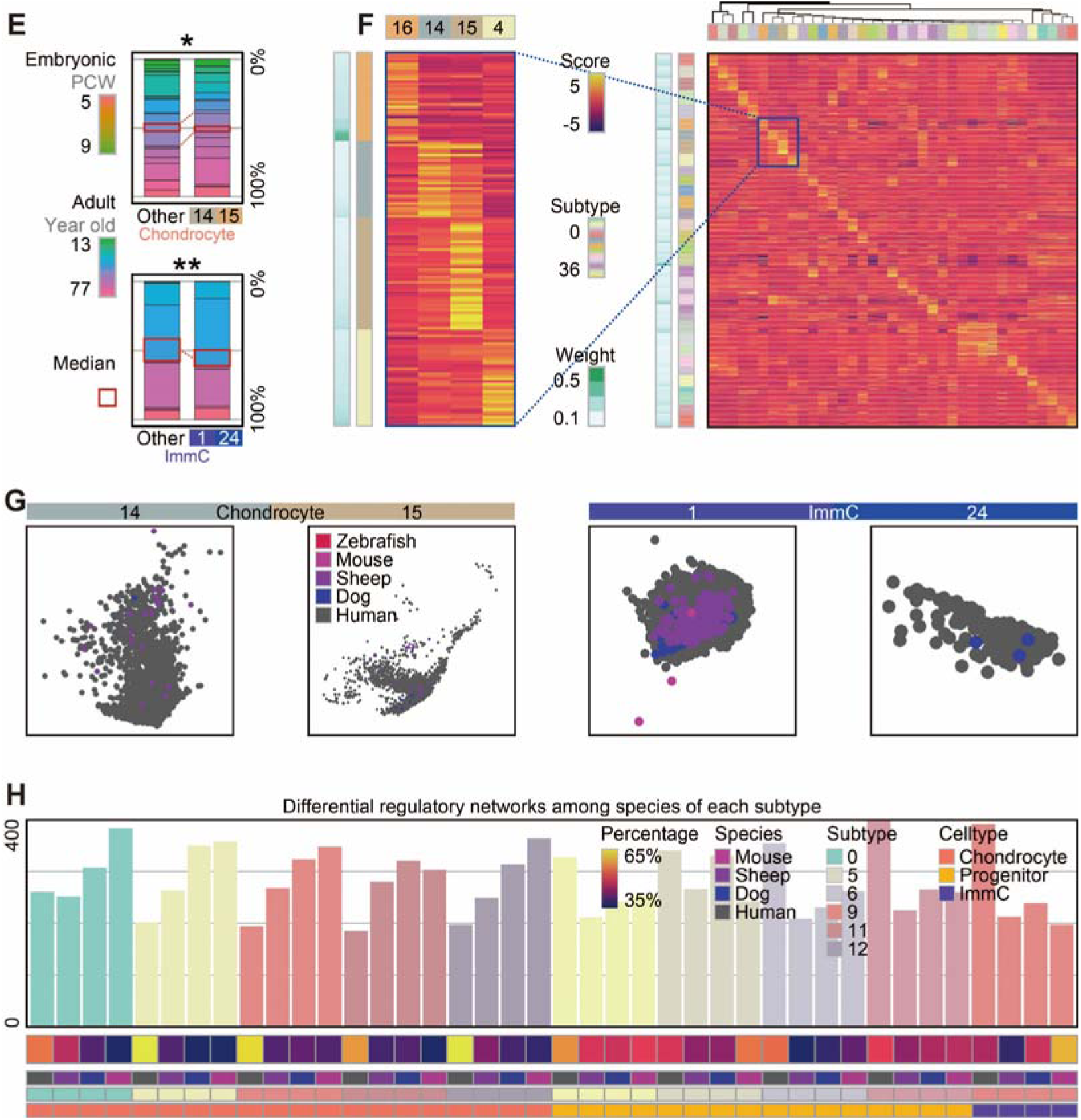
Harmonized annotation of chondrocyte subtypes. (A) Dot plots presenting clustered chondrocytes [21, 24, 37]. Dot color corresponding to clustering resolution and dot size to count number. Arrow transparency corresponding to proportion and arrow color to count number. (B-D) Sankey plots showing the one-to-one relationship of cell identities between JointMap-classified subtypes and available original annotations (Table S1) of chondrocytes (B), progenitors (C), and ImmCs (D). (E) Top histogram showing age information in chondrocyte subtypes and the significant difference (*p* = 0.045) using asymptotic two-sample Fisher-Pitman permutation test [25, 38]. Same test in ImmCs showing significant difference (*p* = 0.007; bottom). * *p* < 0.05; ** *p* < 0.01; *** *p* < 0.001; **** *p* < 0.0001. Abbreviation including human post-conception weeks (PCW). (F) Heatmap presenting the scaled average expression of the top 50 machine learning genes of each chondrocyte subtype. (G) Embeddings showing species of chondrocyte subtype 14/15 and ImmC subtype 1/24. (H) Top histogram showing the numbers of regulons with significantly different activities of each subtype among species (Table S2). Bottom heatmap annotating the percentage of regulons that differ among species at major cell population level (Table S2).

**Figure S3.**
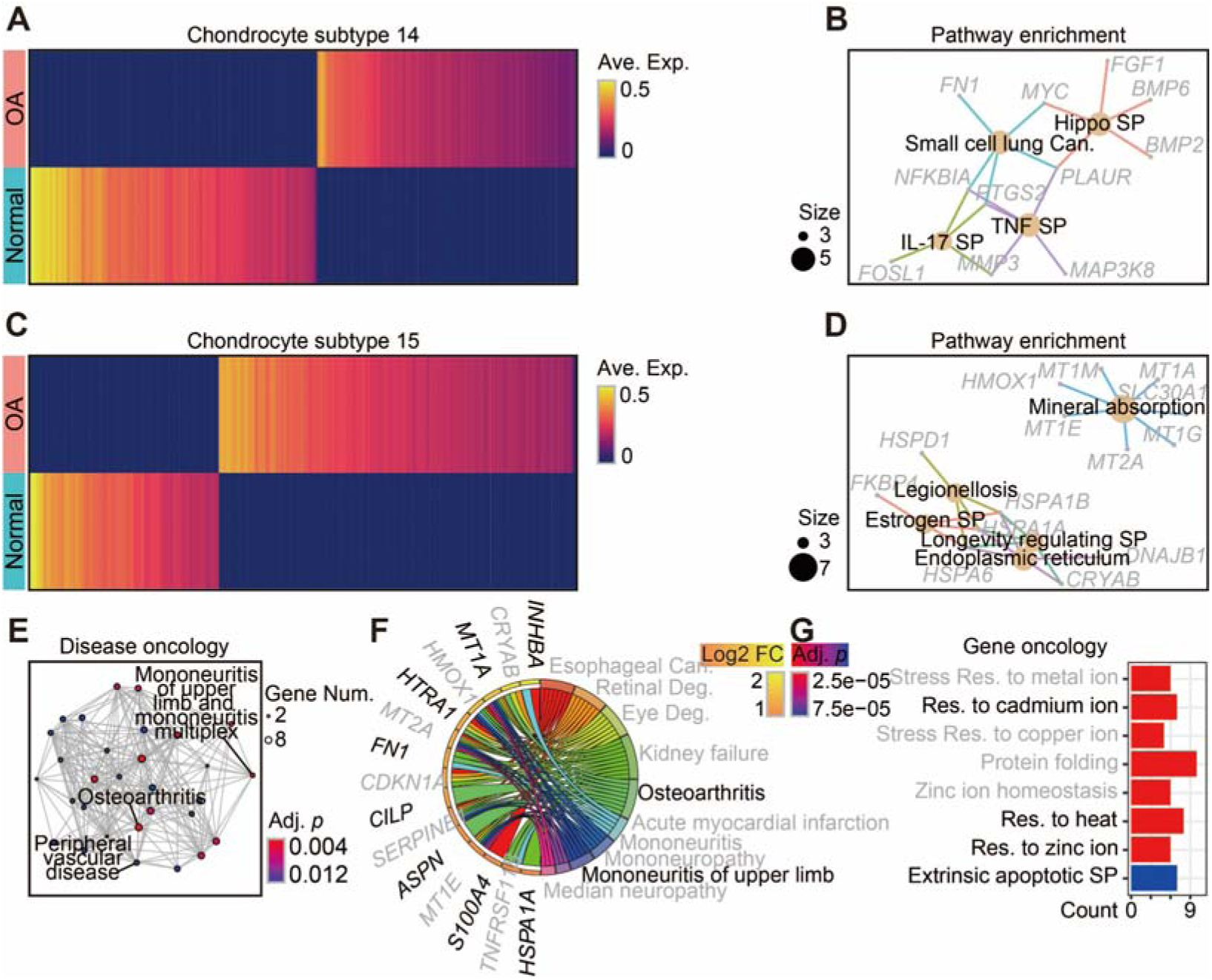
Arthritic transcriptional change in SERPINA3^+^ chondrocytes. (A-B) Heatmap (A) presenting the DEGs in OA and net plots (B) showing the KEGG enrichment analysis in chondrocyte subtype 14. Abbreviation including signaling pathway (SP). (C-G) Enrichment for DEGs in OA (C) through KEGG (D), DO (E-F), and GO (G) enrichment analysis in chondrocyte subtype 15. Abbreviations including degeneration (Deg.) and response (Res.).

**Figure S4.**
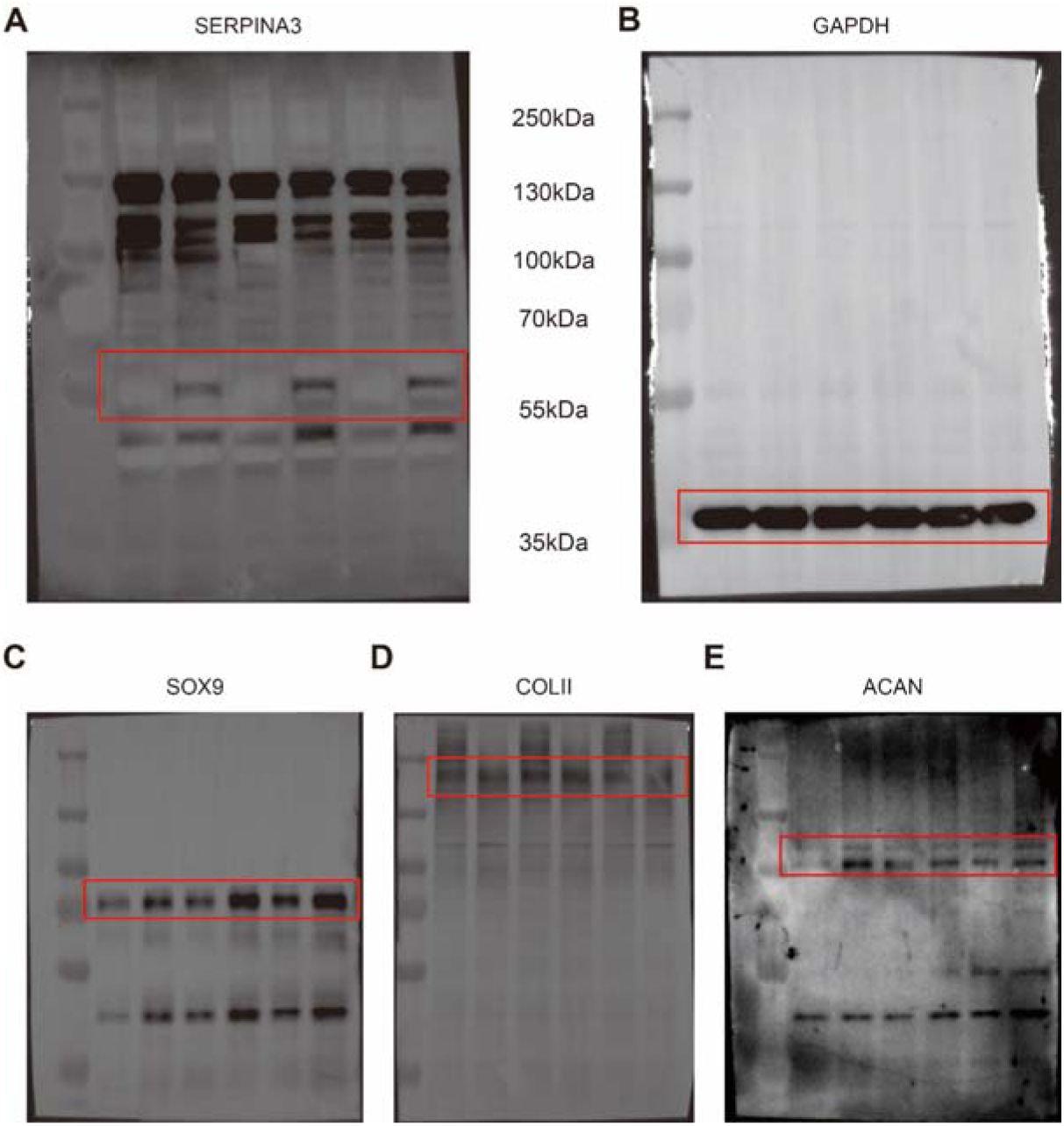
SERPINA3 promotes chondrogenesis in human OA chondrocytes. Uncropped membranes showing the Western blot of SERPINA3 (A), GAPDH (B), SOX9 (C), COLII (D), and ACAN (E).

**Figure S5.**
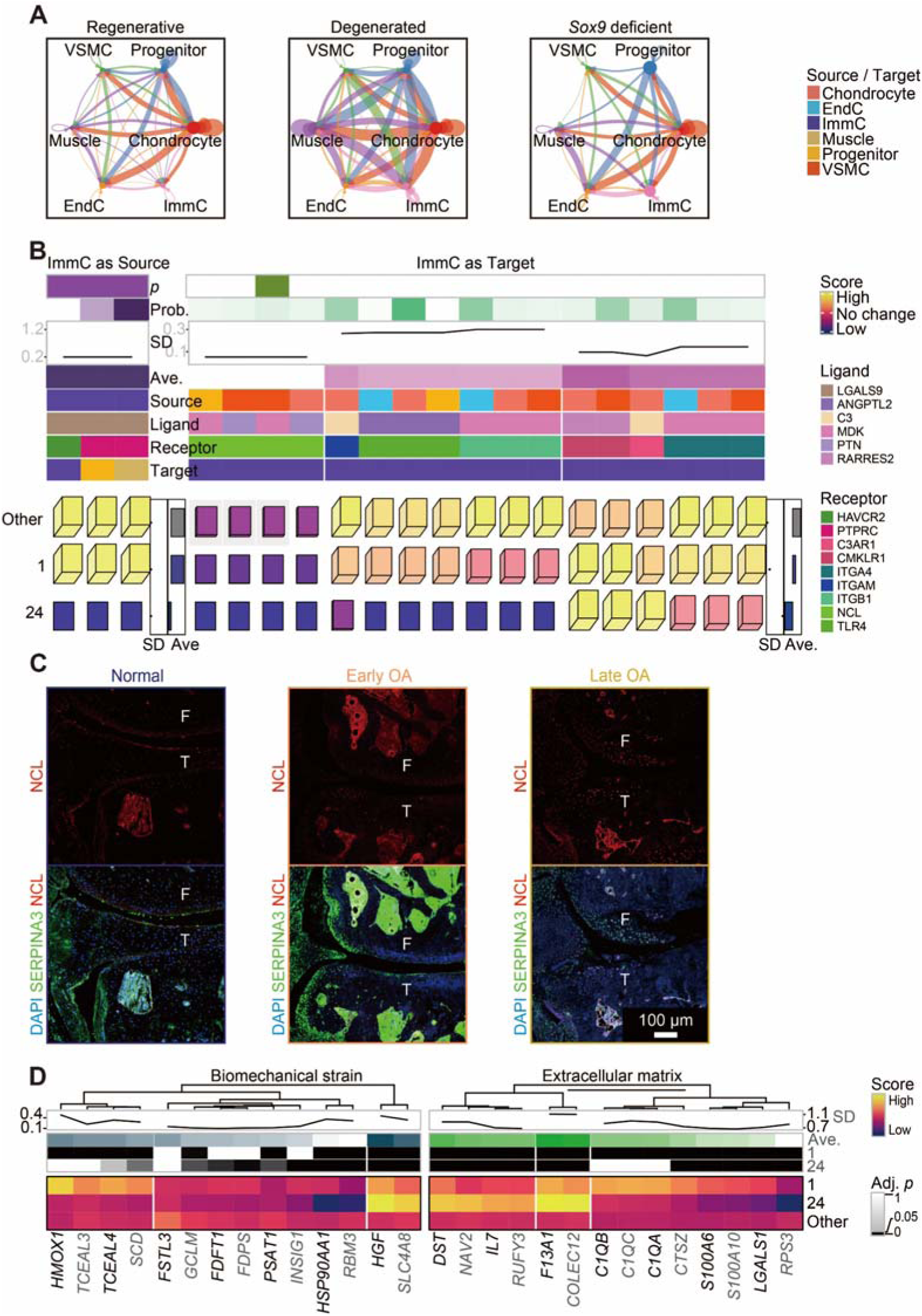
Microenvironment and extracellular matrix. (A) Circular dot plots presenting the crosstalk among cell types of chondral microenvironment under regenerative, degenerated, and *Sox9* deficient conditions speculated by CellChat [49]. (B) Heatmaps presenting the scores (normalized average expression of ImmCs) of genes (ligands in left heatmap or receptors in right heatmap). (C) Anatomical sampling and immunostaining of DAPI (blue), SERPINA3 (green) for chondrocyte subtype 14/15, and NCL (red) in mouse MMT-OA model. (D) Heatmaps presenting the scores (normalized average expression of ImmCs) of genes related to biomechanical stain (GSE165874; left) and ECM from AmiGO (right) [51].

**Figure S6.**
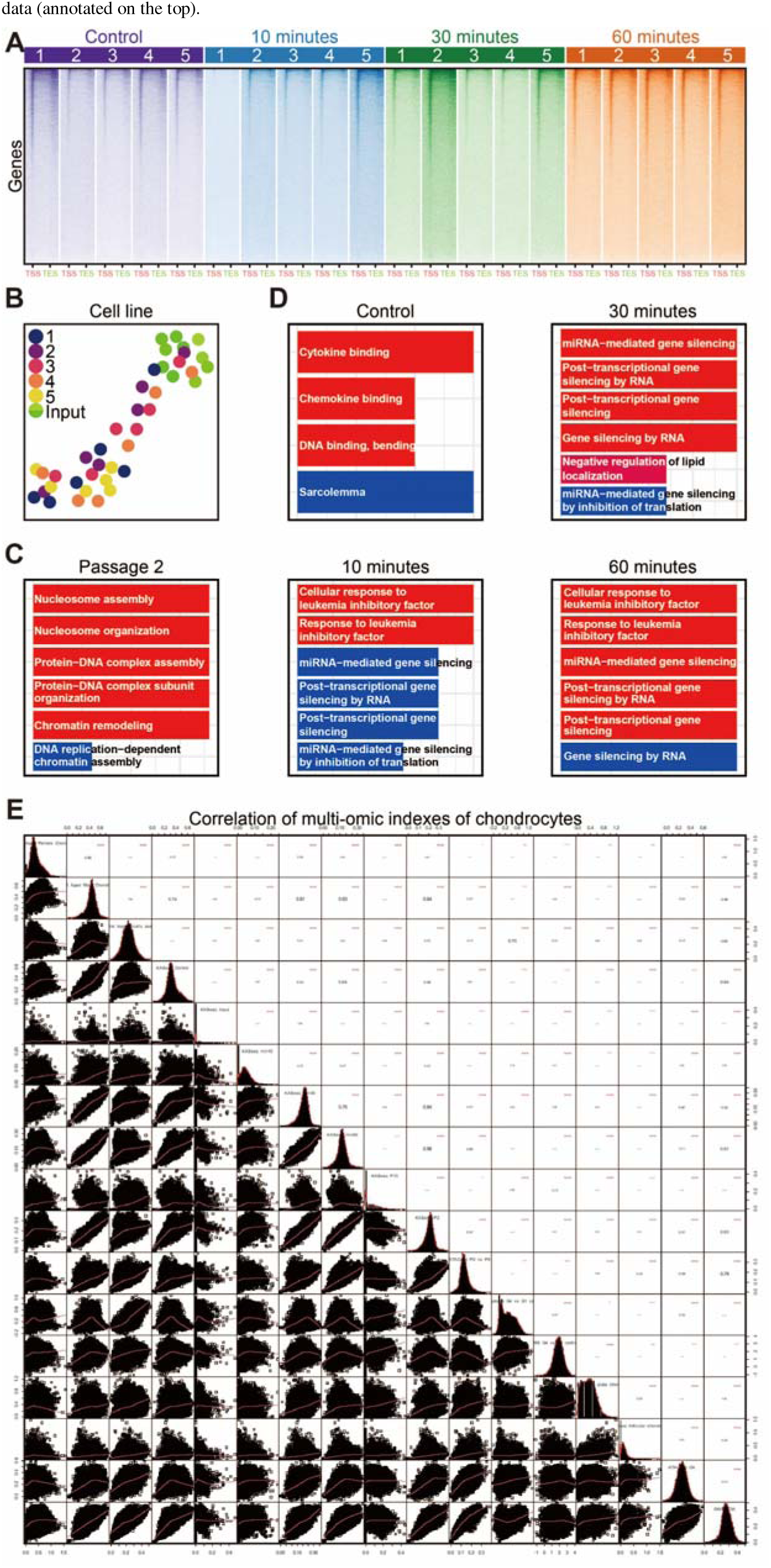
Multi-omic integration of chondrocytes in JointMap. (A) Heatmap showing KAS-seq signal distribution at gene-coding regions under experiments of different times. (B) Embedding showing samples of different cell lines. (C) Bar plots of enrichment results based the genes shown in Fig. 6D of P2. (D) Bar plots of enrichment results based the genes shown in Fig. 6D of control, 10 minutes, 30 minutes, and 60 minutes groups. (E) Dot plots showing the correlation between multi-omic indexes. The names of columns/rows are “LCMS - Aged female/male Chondrocyte”, “ATAC-seq - Chondral TAD”, “ KASseq - control/input/10min/30min/60min”, “ KASseq - P13/P2”, “ATAC-seq - P0 *vs*. P8”, “Trascriptome/LC-MS - G4 *vs*. G1 cartilage”, “ATAC-seq - Growth plate chondrocyte/articular chondrocyte”, “ATAC-seq - OA”, and “GWAS - OA”, respectively.

**Figure S7.**
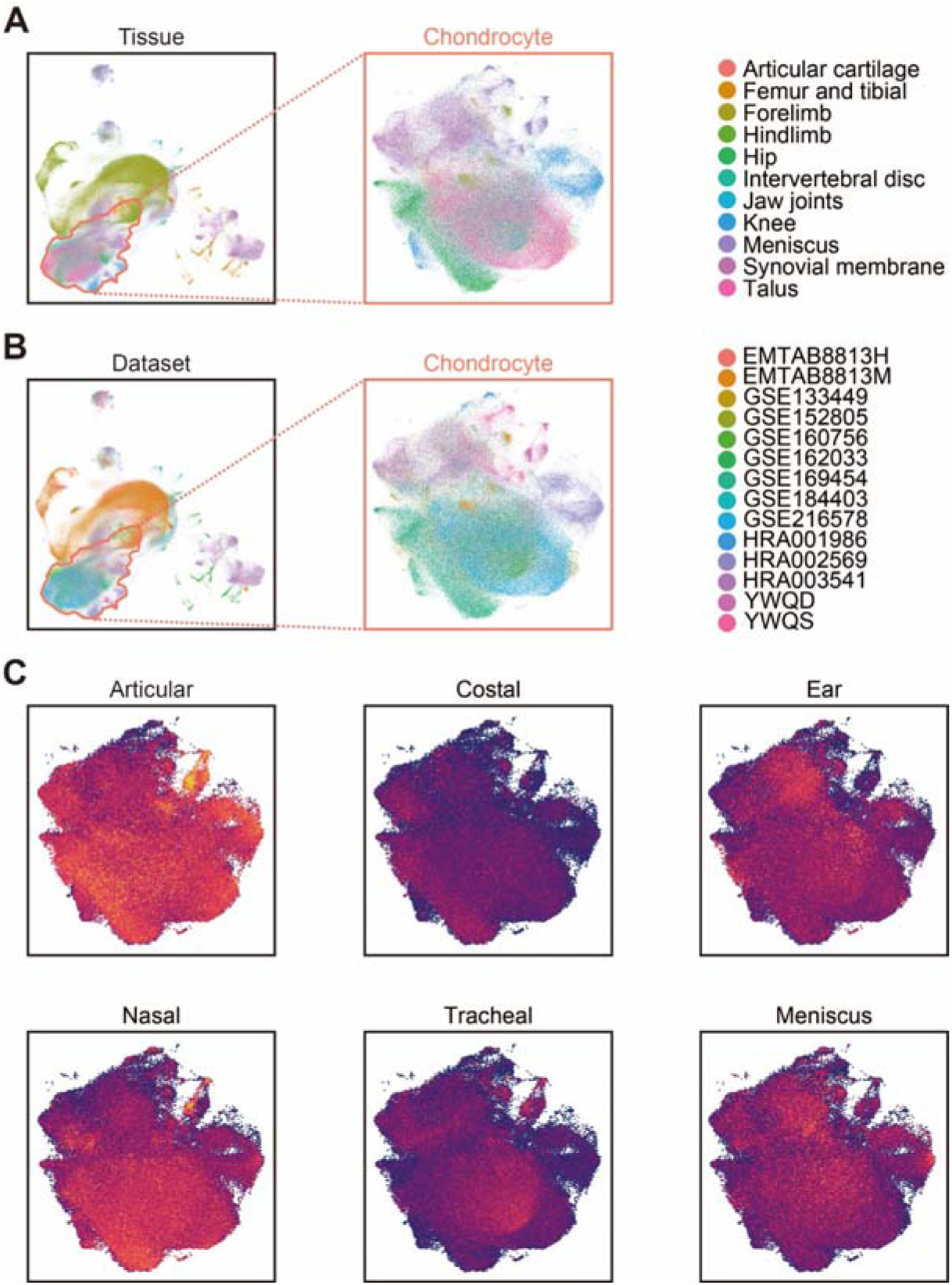
Spatial traits of chondrocyte subtypes. (A) Embedding showing all cells (left) and chondrocytes (right) colored by the original tissues of the sequenced samples. (B) Embedding showing all cells (left) and chondrocytes (right) colored by the sources of datasets (Table S1). (C) Embedding showing the chondrocytes colored by the spatial module scores of diverse pig cartilaginous tissues corresponding to Fig. 7A.

**Figure S8.**
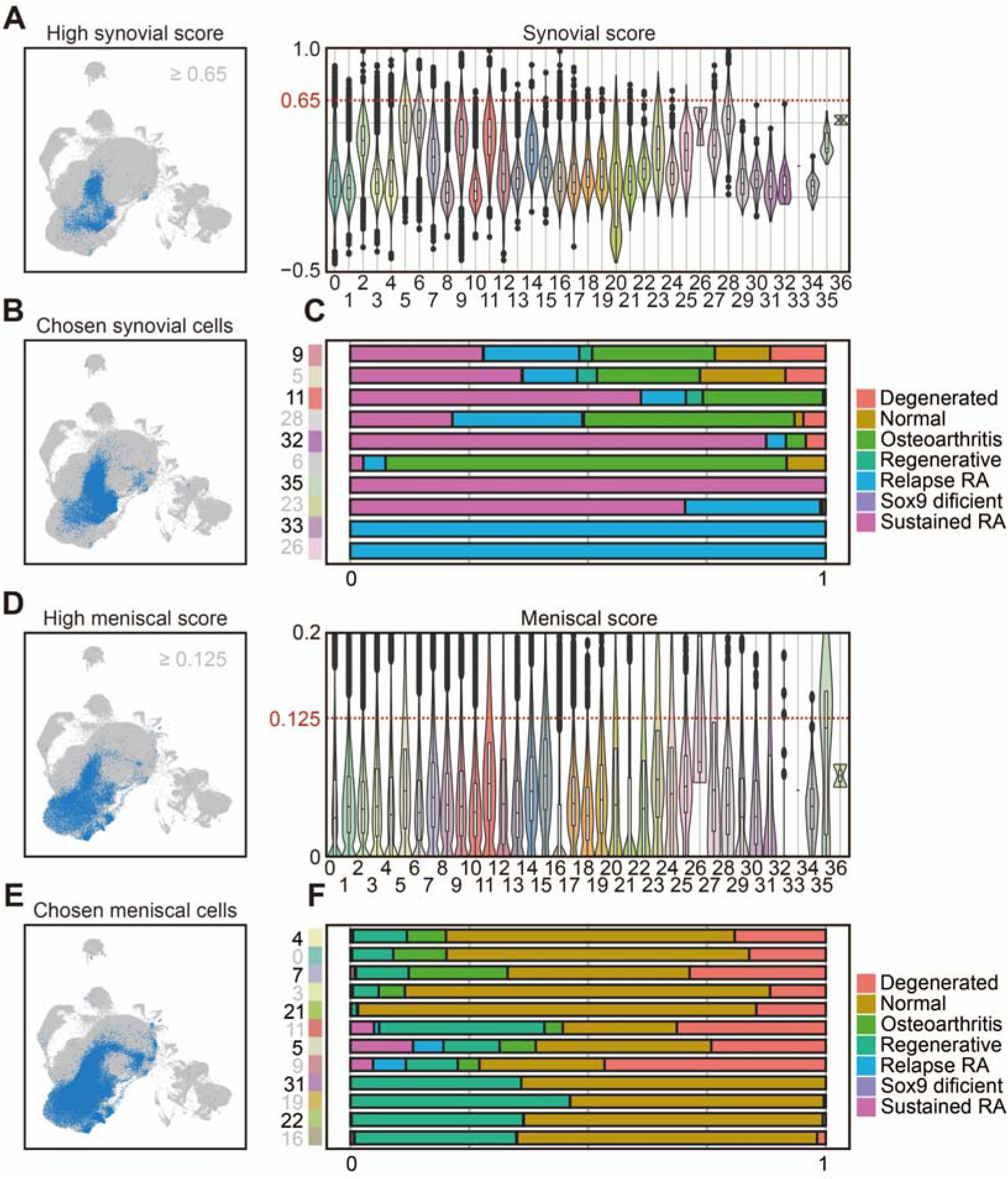
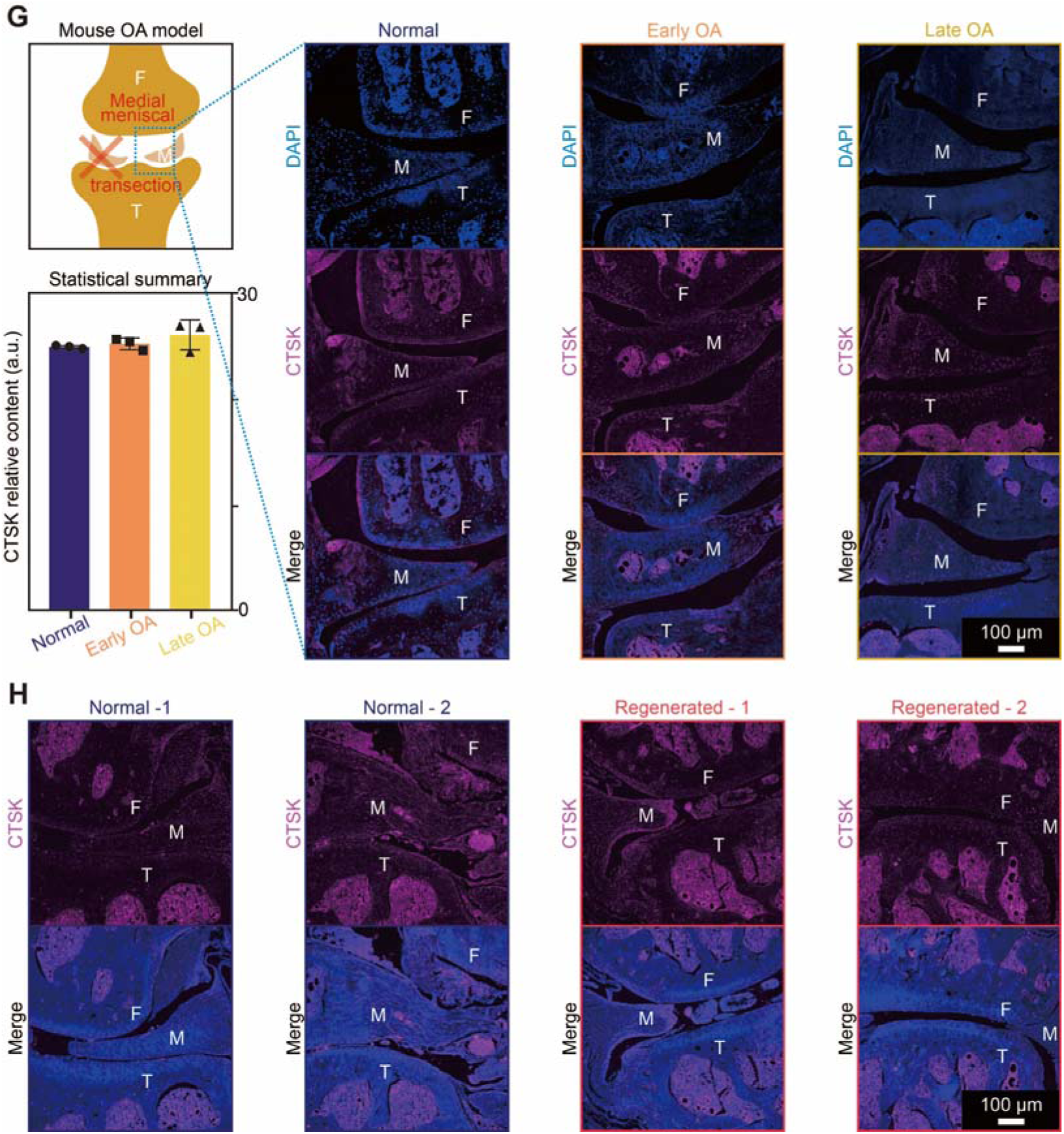
Exploration of synovial and meniscal cells. (A) Embedding showing all cells with synovial scores higher than 0.65 colored in blue. Violin plots showing the synovial scores colored by subtypes. (B) Embedding showing all chosen cells in blue (as showed in Fig. 8A and Fig. S8A) for synovial exploration. (C) Bar plots presenting the compositional percentage of each synoviocyte subtype. Bar color representing the disease conditions of the sequenced samples. (D) Embedding showing all cells with meniscal scores higher than 0.125 colored in blue. Violin plots showing the meniscal spatial module scores calculated based on meniscal specific genes colored by subtypes. (E) Embedding showing all chosen cells in blue (as showed in Fig. 8E and Fig. S8D) for meniscal exploration. (F) Bar plots presenting the compositional percentage of each meniscal chondrocyte subtype. Bar color representing the disease conditions of the sequenced samples. (*continue*). (G) Anatomical sampling and immunostaining of DAPI (blue) and CTSK (red) for meniscal chondrocyte subtype 11 in mouse MMT-OA and control samples. On the left, schematic diagram showing the mouse MMT-OA model, and bar plots demonstrating the protein level of CTSK. Abbreviation including meniscus (M). (H) Anatomical sampling and immunostaining of DAPI (blue) and CTSK (red) for meniscal chondrocyte subtype 11 in the mouse meniscal regeneration model [4, 40].

