## Supplementary text and tables for "JointMap identified SERPINA3^+^ chondrocytes as a therapeutic target for osteoarthritis": Supplementary.docx

**Supplementary legend**

**Table S1.** Sheet 1 annotating collected single-cell datasets, with corresponding species, disease condition, chondral tissue, sample name, and research paper. Sheet 2 showing the cross-omic sources and the detailed indexes.

**Table S2.** Sheet 1 listing reported chondral subtypes and their marker genes. Sheet 2-7 presenting the weighted markers of each subtype speculated by machine learning. Sheet 8-9 showing the interspecies difference in celltype-level and in subtype-level. Sheet 10-13 demonstrating results of enrichment analysis on markers of each chondrocyte subtype.

**Table S3.** Table of DO, GO, and KEGG enrichment analysis based on the DEGs between OA and healthy controls of chondrocyte subtype 14/15.

**Abbreviations**

Adj.: adjusted; ATAC-seq: assay for transposase-accessible chromatin with high throughput sequencing; a.u.: arbitrary unit; Ave.: average; αMEM: α Minimum Essential Medium; BDM: basal differentiation medium; Can.: cancer; DEG: differentially expressed gene; Deg.: degeneration; Dev.: development; Dis.: disease; DO: disease ontology; E: embryonic; ECM: extracellular matrix; EndC: endothelial cell; Exp.: expression; F: femur; FBS: fetal bovine serum; FC: fold-change; G: grade; GO: gene ontology; GSEA: gene set enrichment analysis; hPSC: human embryonic stem cell; ImmC: immune cell; KAS-seq: kethoxal-assisted single-stranded DNA sequencing; kDa: kilodalton; KEGG: Kyoto encyclopedia of genes and genomes; LC-MS: liquid chromatography-mass spectrometry; M: meniscus; MMT: medial meniscal transection; Neg.: negative; Norm.: normal; OA: osteoarthritis; OE: overexpression; P: passage; PCA: principal component analysis; PCW: post-conception week; Pos.: positive; Prob.: probability; qPCR: quantitative polymerase chain reaction; RA: rheumatoid arthritis; Reg.: regulation; Reg-Act: regulons activity; Res.: response; ROI: region of interest; SD: standard deviation; Sig.: signal; SP: signaling pathway; T: tibia; TAD: topologically associating domain; TES: transcription end site; TSS: transcription start site; UMAP: uniform manifold approximation and projection; VSMC: vascular smooth muscle cell
